# Highly multiplexed design of an allosteric transcription factor to sense novel ligands

**DOI:** 10.1101/2024.03.07.583947

**Authors:** Kyle K. Nishikawa, Jackie Chen, Justin F. Acheson, Svetlana V. Harbaugh, Phil Huss, Max Frenkel, Nathan Novy, Hailey R. Sieren, Ella C. Lodewyk, Daniel H. Lee, Jorge L. Chávez, Brian G. Fox, Srivatsan Raman

## Abstract

Allosteric transcription factors (aTF), widely used as biosensors, have proven challenging to design for detecting novel molecules because mutation of ligand-binding residues often disrupts allostery. We developed Sensor-seq, a high-throughput platform to design and identify aTF biosensors that bind to non-native ligands. We screened a library of 17,737 variants of the aTF TtgR, a regulator of a multidrug exporter, against six non-native ligands of diverse chemical structures – four derivatives of the cancer therapeutic tamoxifen, the antimalarial drug quinine, and the opiate analog naltrexone – as well as two native flavonoid ligands, naringenin and phloretin. Sensor-seq identified novel biosensors for each of these ligands with high dynamic range and diverse specificity profiles. The structure of a naltrexone-bound design showed shape-complementary methionine-aromatic interactions driving ligand specificity. To demonstrate practical utility, we developed cell-free detection systems for naltrexone and quinine. Sensor-seq enables rapid, scalable design of new biosensors, overcoming constraints of natural biosensors.

## Introduction

Allosteric proteins pervade biology. They drive virtually all cellular processes as sensors, regulators, enzymes, and signaling proteins. Designing allosteric proteins is key to our efforts to engineer biology. One such class of allosteric proteins is allosteric transcription factors or aTFs. aTFs are universal molecular switches, governing gene expression in response to diverse cellular and environmental cues^1–3^ . These regulators modulate transcription by changing their affinity to specific operator sequences, often upon binding to small molecules. This simple yet powerful genetic control mechanism cements aTFs as foundational elements for small-molecule biosensing in synthetic biology^4–12^.

Despite the versatility of aTFs as molecular switches, a critical limitation in the field is the ability to create aTFs specifically tailored to sense particular target molecules. This shortcoming hinders the precise customization of aTFs for applications such as metabolic engineering, circuit design, and biosensing, where the ability to respond to defined ligands selectively is paramount. We currently rely almost exclusively on natural aTFs, of which only a small number (∼20-25) are well characterized for biosensing applications^4,12–15^. Although bioprospecting has uncovered new aTFs for specific molecules, this strategy is slow and impractical for the discovery of aTFs designed to sense any arbitrary molecule of interest (Fig. 1a)^16,17^. A promising alternative is to engineer the specificity of known aTFs toward new ligands. However, the examples of evolved aTFs with successfully altered specificities still only bind ligands structurally similar to the wildtype ligand, limiting the overall utility of directed evolution to design biosensors for entirely new chemical classes^13,18–20^.

**Figure 1.**
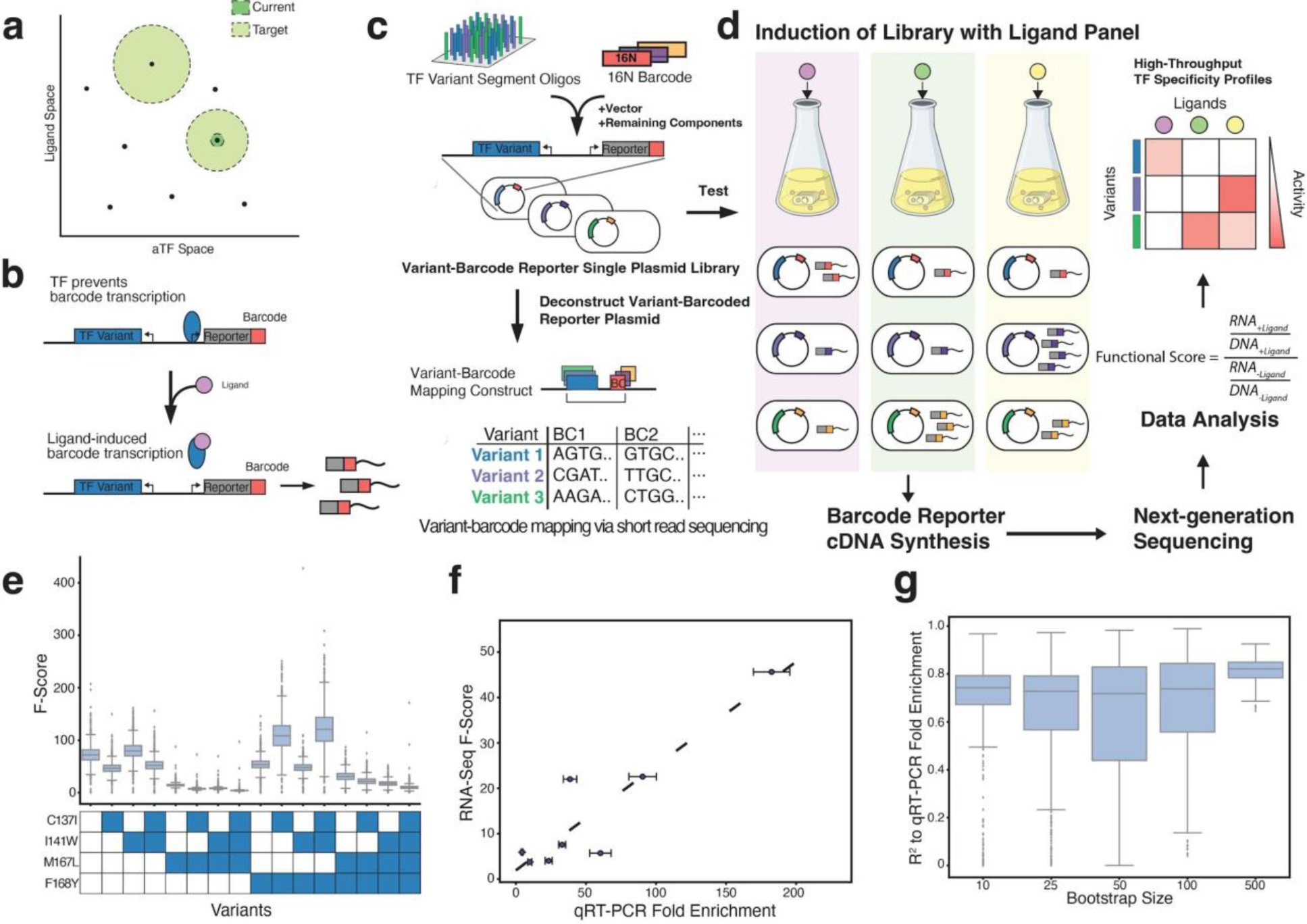
An RNA-Seq approach for high-throughput aTF characterization. ***(a)*** The space of transcription factors and small molecules that can be sensed with different approaches. Black points represent specific ligand:aTF pairs. The dark green circle represents the extent to which existing methods can currently expand aTF:ligand affinity. The light green circle represents the extent to which new methodologies must increase aTF:ligand pairs. ***(b)*** Construct design pairs aTF variant activity to transcription of randomized barcodes. ***(c)*** Construction and mapping of libraries to barcodes. Gene segments of aTFs with mutations are synthesized in oligo pools. Single-plasmid libraries are constructed with oligos, random 16N barcodes, plasmid vector, and remaining components of the target gene. These constructs are deconstructed for short-read mapping. ***(d)*** Original plasmid library is tested under different ligand conditions, resulting RNA is sequenced, and functional scores (F-scores) are generated to obtain aTF specificity profiles. ***(e)*** Box plots of fold enrichment for each TtgR variant via RNA-Seq. The box represents the interquartile range. Whiskers extend to 1.5 times the IQR. Fliers denote points that lie outside the whiskers. Filled blue square indicates the presence of mutation in the variant. ***(f)*** Correlation of qRT-PCR data and fold enrichment from RNA-Seq. qRT-PCR fold enrichment was measured via biological replicates of clonal strains of 8 of the 16 variants (see Methods). The RNA-Seq fold enrichment value was calculated by summing the counts of all barcodes associated with a particular variant (see Methods). The R^2^ for this dataset is 0.83. ***(g)*** Bootstrap correlation of qRT-PCR fold enrichment to RNA-Seq data for 8 of the 16 variants. Groups of 10, 25, 50, 100, or 500 barcodes were sampled for each variant across 500 cycles. The resulting correlation for each cycle is plotted.

Altering the ligand specificity of allosteric proteins presents a formidable challenge in library design and screening. The challenge arises from the tight interconnection of residues involved in ligand binding with those crucial for allosteric actuation^21–24^. Therefore, libraries designed solely to optimize protein-ligand interactions, as one might for designing binders, without considering the role of residues in allostery, are likely to abrogate the switch-like properties of aTFs. Furthermore, successful designs within a library are typically rare and may display weak activity toward non-native ligands. We need a high-throughput screening method with high sensitivity to effectively identify these rare, low-activity ‘hits’ amid a vast pool of non-functional designs. These low-activity variants are useful scaffolds from which high-activity variants can ultimately be derived by iterative design or directed evolution. Additionally, data from low-activity variants, which are commonly overlooked, can be leveraged to strengthen machine-learning models. Unfortunately, conventional enrichment techniques (like flow cytometry, plate-based screening, or growth selections) cannot identify these important variants owing to their lack of sensitivity, scale, or both^13,25–29^.

We report Sensor-seq, a platform for creating aTF biosensors with high sensitivity and scale (Fig. 1b). Sensor-seq combines phylogeny-guided sequence diversification for library design with an RNA barcoding system to screen aTF variants through deep sequencing. As a starting scaffold for design, we sought a promiscuous aTF with a large binding pocket that could accommodate diverse ligands. We reasoned that designed mutations could customize the binding pocket with shape complementary interactions for a target ligand. Our rationale is supported by prior research demonstrating that enzyme evolvability and acquisition of novel functions benefit from a promiscuous starting point^31,32^. Therefore, we chose the aTF TtgR, a multi-drug efflux regulator, as our starting scaffold because it is known to bind to several antibacterial molecules^30^. We chose as targets for the design seven non-native ligands of diverse chemical structures – four derivatives of the cancer therapeutic tamoxifen, the antimalarial drug quinine, the opiate analog naltrexone, and the plant secondary metabolite ellagic acid – as well as two native flavonoid ligands, naringenin and phloretin. Six out of the seven non-native molecules are synthetic chemicals for which traditional genome-mining for natural biosensors would likely be ineffective.

We evaluated 17,737 TtgR variants against the panel of seven non-native and two native ligands. Sensor-seq identified novel biosensors for all but one of these ligands, with high dynamic range and diverse specificity profiles. Moreover, our comprehensive dataset allowed us to identify distinct linchpin positions and mutations driving specificity toward each ligand and unravel the molecular principles governing protein-ligand specificities for this scaffold protein. We obtained a crystal structure of a TtgR design bound to naltrexone, a non-native ligand, to elucidate the structural basis of the customizable specificity of TtgR to many ligands, a generally uncommon property among proteins. To illustrate the practical application of these engineered aTFs, we constructed cell-free biosensing systems for detecting naltrexone and quinine, which have potential applications in detecting opioid overdose and wastewater contamination, respectively. In summary, Sensor-seq advances the on-demand design of biosensors for any target molecule. By broadening specificities to include non-native ligands, Sensor-seq liberates us from the restrictions imposed by solely relying on natural biosensors. Finally, the Sensor-seq workflow is adaptable to other proteins whose function can be linked to transcription.

## Results

### Validation of Sensor-seq on a pilot library

Sensor-seq assesses a pooled library of aTF variants by linking their ligand-induced responses to deep sequencing. Each cell expresses a unique aTF variant under a constitutive promoter regulating a reporter locus controlled by the aTF’s native promoter (Figure 1b). The activity of each aTF variant is quantified by measuring reporter transcript RNA levels through RNA sequencing (RNA-seq). The F-score, a normalized ratio of reporter transcript levels in the presence and absence of the ligand, is a quantitative measure of each variant’s activity. The advantage of this approach is that aTF variants that are allosterically inactive either in the constitutively OFF (always repressing transcription) or constitutively ON state (unable to repress transcription) have an F-score of ∼1. These unproductive variants can be disregarded, allowing us to focus on the rare but potentially productive members of the library (F-score > 1).

A key challenge for high-throughput screens of proteins that regulate gene expression in *trans* (like aTFs) is the association of each variant’s genotype with its transcriptional output. Linking the aTF variant to its function (e.g. reporter abundance) presents a technical challenge for mapping genotype to phenotype. We addressed this technical hurdle with a novel barcoding method. Each aTF variant is placed in *cis* with randomized barcodes with an intervening constant region comprising the unmutated part of the aTF and the promoter regulating the reporter (Fig 1b). The randomized barcode is analogous to that in reporters used in most massively parallel reporter assays. However, sequencing the reporter alone would be insufficient for identifying which aTF variant was responsible for transcribing each reporter. To map the aTF to the transcript barcode, the intervening constant region is removed by restriction digest, which places the aTF variant in close proximity to the reporter barcode. The variant and the associated barcode are mapped by traditional short-read deep sequencing (Fig. 1c). The transcript levels of each barcode of this screening construct reveal whether the associated aTF variant responds to the target ligand. Since the barcode region is extremely short, we use high-volume, short-read sequencing to accurately quantify the activity levels of each variant (Fig. 1d). This platform, Sensor-seq, is generalizable and easily repeated to profile the variant library among any number of environmental perturbations (e.g. ligands). Importantly, our mapping scheme scales as a constant with the number of ligands tested. That is, only a single round of mapping is needed for users to test any number of ligands against thousands of aTF variants. *E. coli* containing the plasmid library are dosed with either the target ligand or a vehicle control and harvested in log phase to obtain both total RNA (sequenced as cDNA) and the library plasmid DNA (Fig. 1d). The cDNA count provides a measure of function while the plasmid DNA count is used for normalization. These counts are used to calculate functional scores (F-scores) for each variant induced with different ligands. The library can be incubated with several ligands independently and evaluated in a single pooled sequencing to achieve scale.

To verify the effectiveness of our proposed methodology, we applied Sensor-seq to a small library of 16 TtgR variants previously characterized for their responses to a native ligand, naringenin, using a GFP reporter protein^33^. Gene fragments encoding these variants were incorporated with random 16N reporter barcodes into our screening construct and mapped through short-read sequencing, with each variant mapping to approximately 8,000 barcodes (Supplementary Fig. 1). The F-score of each variant in response to 1mM naringenin was determined using pooled RNA-seq, calculating cumulative barcode counts of the reporter relative to the vehicle control after normalizing to plasmid DNA (See methods) (Fig. 1e). From the 16 variants, 8 spanning the full range of naringenin responses were selected for clonal quantification through qRT-PCR. Comparison of the qRT-PCR fold enrichment and the Sensor-Seq (RNA-seq based) F-score showed a high correlation (R^2^=0.83) (Fig. 1f). In summary, Sensor-seq accurately reproduced activity differences observed via qRT-PCR for a small library.

When dealing with larger protein libraries, the requirement of 8,000 barcodes per variant for sequencing becomes impractical. To evaluate the sensitivity of our F-score measurements to changes in the number of barcodes per variant, we down-sampled the number of barcodes per variant. Using a Monte Carlo sampling approach, we randomly selected 10, 25, 50, 100, and 500 barcodes per variant for 500 trials each and scored each sample based on its correlation to the qRT-PCR dataset (Fig. 1g). Each bootstrap group exhibited, on average, similar correlation to the qRT-PCR assay compared to the 8,000 barcodes per variant. Therefore, Sensor-seq can accommodate larger protein libraries with reduced barcode-to-variant ratio and lower total sequencing reads per variant, minimizing sequencing requirements.

### Identifying novel sensors for non-native ligands

Having validated Sensor-seq, we next sought to apply this approach to redesign the specificity of TtgR toward non-native ligands. We used FuncLib to generate a library of TtgR variants in a ligand-agnostic manner (i.e., not targeted toward any particular ligand) with mutations around the binding pocket. FuncLib combines evolutionary phylogeny of a protein family for diversity generation together with Rosetta calculation of mutational stability for curation to create a library of variants^34–37^. FuncLib delineates a sequence space of allowable point mutations within the ligand-binding site of TtgR by filtering out mutations that are unlikely to occur naturally as determined by a multiple sequence alignment of TtgR homologs. This important filtering step eliminates mutations likely to disrupt allostery but preserves mutations that provide diversity^34–37^. Mutations passing the evolutionary curation test are computationally modeled with Rosetta into the TtgR structure to assess their stability. Only those that maintain protein stability are considered viable point mutations. Based on a list of allowable point mutations using FuncLib, we created a combinatorial library of 17,737 variants containing between 1-4 mutations per sequence.

We selected a panel of seven non-native ligands for which to develop new biosensors and two native ligands to create biosensors with altered specificity toward a native ligand (Fig. 2a). We selected four derivatives of tamoxifen (Tam), a breast cancer therapeutic, to create specific and multi-specific sensors. Tamoxifen is a selective estrogen receptor-modulating prodrug that is converted into active metabolites, 4-hydroxy-tamoxifen (4Hy) and N-desmethyltamoxifen (Ndes). These two metabolites are then catabolized to endoxifen (End)^38,39^. These four metabolites share a core structure with three aromatic rings, one linked to a methylamine sidechain, but they differ in the derivatization of aromatic rings and/or the sidechain. We chose these four molecules to identify biosensors capable of distinguishing among them. Additionally, we selected quinine (Quin), naltrexone (Nal), and ellagic acid (EllA) as non-native ligand targets. Quinine, a malaria therapeutic, features a quinoline ring with two aromatic benzene rings, resulting in a distinctive non-planar structure^40^. Naltrexone, an opioid analog used in addiction treatment, has a tetracyclic arrangement with three fused six-membered carbon rings^41^. Ellagic acid is a plant polyphenol with a planar, symmetric arrangement. (Fig. 2a)^42^. We chose two native ligands of TtgR belonging to the flavonoid family, naringenin and phloretin, with the goal of identifying biosensors that could discriminate between the two molecules^30^. All seven non-native molecules differ significantly from the native flavonoid ligands (Fig. 2a). Since our primary goal was to identify gain-of-function variants for non-native ligands, we did not design against the native ligands, i.e. we aimed to expand TtgR’s ligand-binding repertoire without consideration for native specificity. We reasoned that the potential applications of novel biosensors would not likely involve the native ligands, which are uncommon plant metabolites.

**Figure 2.**
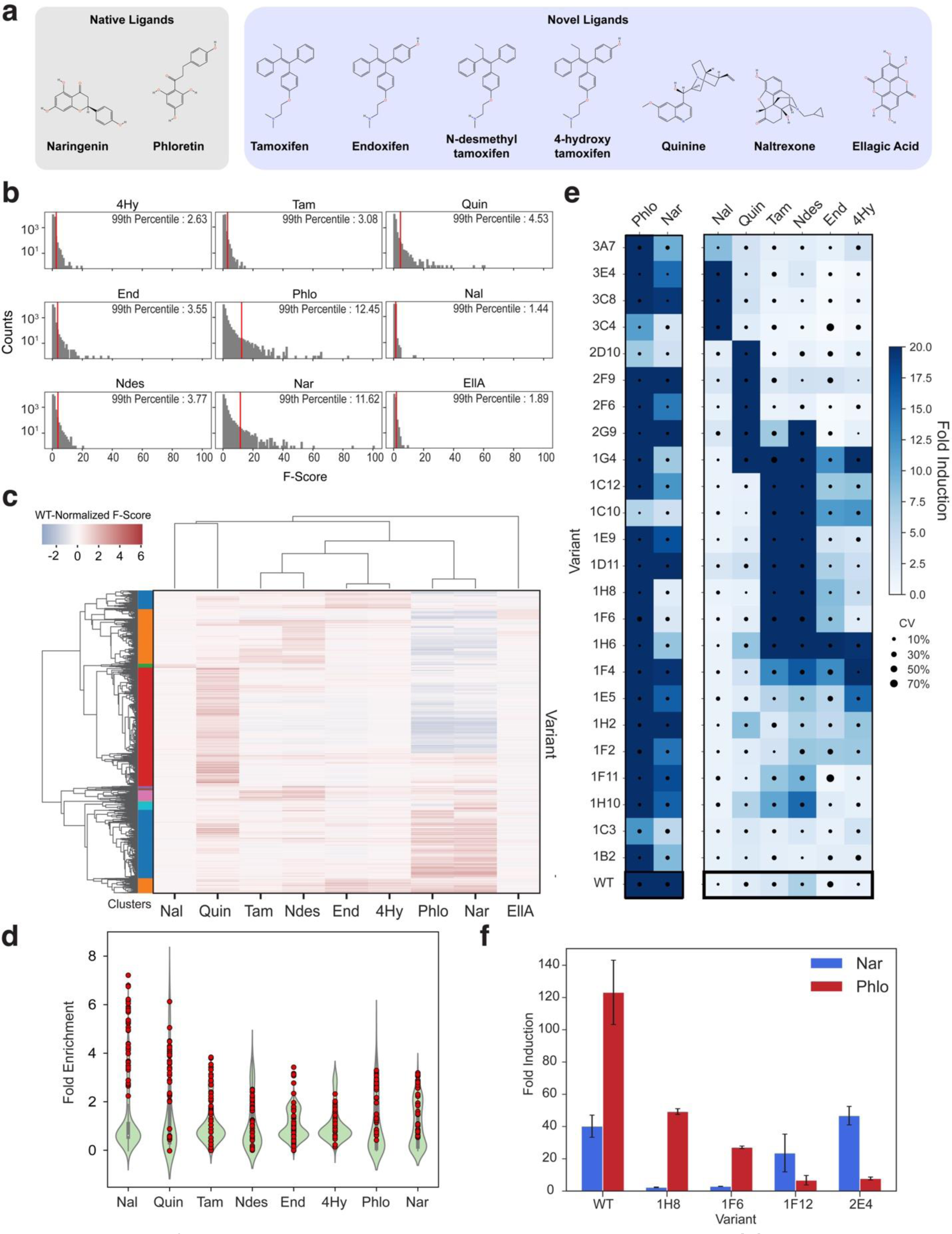
Identifying novel sensors in a ligand agnostic library. ***(a)*** Ligands used in this study. ***(b)*** Histograms showing distribution of F-scores for each variant in the library with each tested ligand. Red line denotes the 99^th^ percentile of F-scores ***(c)*** RNA-Seq fold enrichment data for 16,191 variants which passed filters across nine ligands. Ligands and variants have been clustered via the UPGMA algorithm with a correlation distance metric and a target of 12 clusters (see Methods). The different clusters are denoted by the colored bars on the left of the heatmap. aTF function is shown as the log_2_(F-score) normalized to wildtype. ***(d)*** Violin plot showing distribution of fold enrichment scores of a 251-member library consisting of the top 40 variants for each ligand (determined by F-scores) after fluorescence-based sorting. Fold enrichment scores were calculated as abundance of each library member after induction divided by abundance before induction. Red circles indicate scores of variants selected as top 40 for corresponding ligands. ***(e)*** Heatmap showing fold induction of wild-type TtgR and variants that were selected as top 3 for each ligand based on sorting in (c). Fold induction is calculated as the average of the median fluorescence after induction divided by median fluorescence before induction for three replicates. Right panel consists of non-native ligands, and the left panel consists of the two native ligands. A score ceiling of 20 was imposed for visualization. Black circles within heatmap cells reflect the coefficient of variation which is the standard deviation divided by the mean for each ligand-variant test. ***(f)*** Bar plot of variants with native ligand specificity switches. Fold induction is calculated the same as (d). Error bars represent standard deviations of three replicates.

From a synthesized and cloned variant library of 17,737 members, we observed an average of 20 barcodes per variant for 17,533 variants (98.8%) (Supplementary Fig. 2). We removed variants with a high degree of noise based on the coefficient of variation across biological replicates (see methods). Variants that did not meet this threshold were excluded from further analysis. Reporter RNA was quantified in cells containing the library, which were incubated with each of the nine ligands or their respective vehicle controls (ethanol, DMSO, or water). We calculated the distribution of F-scores for each ligand to quantitatively assess the library’s performance (Fig. 2b, Supplementary Fig. 3, Supplementary Table 5). The F-score at the 99^th^ percentile provided an estimate of the activity and prevalence of variants responding to a particular ligand. Naringenin and phloretin exhibited the highest F-scores at the 99^th^ percentile (11.6 and 12.5, respectively), followed by quinine (4.5), N-desmethyltamoxifen (3.8), endoxifen (3.6), tamoxifen (3.1), and 4-hydroxytamoxifen (2.6). In contrast, both naltrexone and ellagic acid had the lowest F-scores (1.4 and 1.9) (Fig. 2b), suggesting that hits for these ligands, if present in the library, were exceedingly rare. As a reference, the wildtype TtgR’s F-scores for its native ligands phloretin (2.0) and naringenin (1.9) were considerably lower than those observed for the library at the 99^th^ percentile on the non-native target ligands (Supplementary Fig. 4). This suggests that hits for non-native ligands were likely real.

We performed unsupervised agglomerative clustering of 16,191 filtered variants against the nine ligands based on their functional profiles across all ligands (vertical) and the chemical relatedness of ligands (horizontal) with the activity scaled relative to wildtype TtgR (Fig. 2c, Supplemental Fig. 5). Activity on naringenin and phloretin (native ligands of TtgR) showed that nearly half the FuncLib-derived library retained ligand-induced allosteric response on par with or exceeding that of wildtype TtgR. Moreover, ∼85% of TtgR variants demonstrated the ability to repress transcription (Supplementary Fig. 6). As a point of reference, the percentage of repression-competent variants was as low as 15% in our previous LacI variant library which was designed without FuncLib^13^. These results collectively suggest that FuncLib adeptly navigated the sequence space to preserve a high fraction of allosterically active and repression-competent variants in the library, thereby increasing the likelihood of successful designs.

Most TtgR variants showed concordant activities on the native ligands, naringenin and phloretin, similar to wildtype TtgR (Supplementary Fig. 7). However, a small fraction exhibited differential activities toward one molecule or the other– either higher activity on naringenin than phloretin or vice versa. These included variants that lost activity on one ligand but maintained wildtype-like activity on the other (Fig. 2c, blue and white shades), and those that retained wildtype-like activity on one and outperformed wildtype on the other (Fig. 2c, white and red shades). We also observed differential activities among the four tamoxifen derivatives (Fig. 2c). Notably, there was minimal overlap between variants responding to the pair of hydroxylated derivatives, 4-hydroxytamoxifen and endoxifen, and the pair of non-hydroxylated derivatives, tamoxifen and N-desmethyl-tamoxifen. Further, a small group of variants could discriminate between tamoxifen and N-desmethyl-tamoxifen. A substantial fraction of sequences responded to quinine and these quinine-responsive sequences constituted an independent group that did not appear to overlap with variants responding to other ligands (Fig. 2c). A few variants appeared to respond to naltrexone and ellagic acid, albeit weakly in the case of ellagic acid (Fig. 2c). These results show that the Sensor-seq can identify allosterically responsive variants to non-native inducers and successfully distinguish structurally similar ligands to create specific aTF biosensors.

We carried out secondary and tertiary screens to validate the hits from the primary Sensor-seq screen. The secondary screen involved pooling the top 40 variants for each ligand in a cell-based reporter fluorescence assay, and the tertiary screen included clonally assaying individual variants from the secondary screen. For the secondary screen, we created a mini-library by resynthesizing 251 variants corresponding to the top performing variants for each ligand, including overlapping hits between different ligands (e.g. naringenin vs. phloretin or tamoxifen derivatives) (Supplementary Fig. 8, Supplementary Table 6). From this mini-library, we sorted variants capable of repressing GFP expression in the absence of any small molecule. (Supplementary Fig. 9-10). These repression-competent variants were then exposed to each ligand and the high fluorescence cells were isolated and sequenced (Supplementary Fig. 9 and 11). Each variant was scored based on the percentage fold change in abundance between the high fluorescence and repressed sorted populations. The secondary screen was consistent with the results of the primary Sensor-seq results (Fig. 2d, Supplementary Table 7). The top-performing variants for each ligand, as determined by F-scores obtained through Sensor-seq (indicated by red dots), consistently exhibited high fold enrichment when evaluated using the GFP reporter assay (Fig. 2d). However, certain variants associated with N-desmethyltamoxifen, naringenin, phloretin, and quinine gave considerably lower fold enrichment changes in the GFP assay compared to their Sensor-seq scores. This observation raises the possibility that the primary Sensor-seq screen is susceptible to false positives. Naltrexone showed the most consistent response between the two assays in the top 40 variants. We found no hits for ellagic acid in the GFP assay, and thus this ligand was excluded from further analysis. As a tertiary screen, we clonally tested the top three unique variants for each ligand using the GFP reporter by measuring the fold induction: the ratio of mean cell fluorescence in the presence of the ligand and in the absence of the ligand (see Methods). As expected, WT TtgR had a strong response to native ligands, phloretin and naringenin, with only weak activity on N-desmethyltamoxifen and little to no activity on the other non-native ligands (Fig. 2e).

We observed remarkable diversity in the specificity profiles of different variants (Fig. 2e). Variants that acquired specificity for naltrexone (3A7, 3E4, 3C8, and 3C4) showed varied responses to the ligand, ranging from 9- to 43-fold induction. These monospecific variants did not respond to the other non-native ligands. Mono-specificity was also observed with 3 (2D10, 2F9, and 2F6) of the 5 strong quinine responders (Fig. 2e). In particular, one variant, 2D10, exhibited mono-specificity toward quinine with minimal activation from even the native ligands. The two other quinine-responsive variants showed varying degrees of broader specificity. Specifically, 2G9 was additionally activated by tamoxifen and N-desmethyltamoxifen, while 1G4 was activated by all non-native ligands except naltrexone, with fold induction ranging from 13- (endoxifen) to 98-fold induction (tamoxifen). Among variants with broad specificity, the activities toward each ligand varied considerably. For instance, 1H6 shared a similar specificity profile as 1G4, except this variant had high activity toward endoxifen (21-fold induction) at the cost of reduced quinine response (8-fold induction) (Fig. 2d, Supplementary Table 8). 1F4 is another variant that shared a similar profile to 1G4 and 1H6 but showed high activity on 4-hydroxytamoxifen (29-fold induction) and a more muted response to endoxifen, N-desmethyltamoxifen, tamoxifen, and quinine (Fig. 2e, Supplementary Table 8). Eight variants (1C12, 1C10, 1E9, 1D11, 1H8, 1F6, 1F11, and 1H10) appeared to be largely bi-specific to N-desmethyltamoxifen and tamoxifen (both methylated), with activities ranging from 8- to 90-fold induction (Fig. 2e, Supplementary Table 8). However, 1C12 and 1C10 displayed weak to moderate activity on the other two tamoxifen derivatives, 4-hydroxytamoxifen and endoxifen (both hydroxylated) (Fig. 2e, Supplementary Table 8). We did, however, find variants specific to only 4-hydroxytamoxifen and endoxifen in the secondary screen.

Most variants that had high activity on other ligands retained activity on naringenin and/or phloretin, which is consistent with our design criterion of not selecting against native ligand responders. Our data also suggest true specificity switches away from native function may be difficult because binding pockets that could accommodate the larger non-native ligands can also fit the native ligands. (Fig. 2e. Supplementary Table 8). Notably, 2F9 had an even greater response to naringenin and phloretin (101 and 234-fold, respectively) compared to wildtype TtgR (40 and 123-fold, respectively). Two variants (1H8 and 1F6) showed specificity with dramatically higher activity on phloretin over naringenin (49-vs. 2-, and 27-vs. 3-fold induction, respectively). To determine if specificity for naringenin can be favored over phloretin, we clonally evaluated two variants (1F12 and 2E4), which showed a preference for naringenin from our secondary screen. 1F12 and 2E4 retained near-WT responses for naringenin (24- and 47-fold induction, respectively) with a 15- and 18-fold loss in activation with phloretin compared to WT (7- and 8-fold induction, respectively) (Fig. 2f).

We next evaluated the predictive performance of Sensor-seq using the results from the tertiary screen as “ground truth”. We observed a strong relationship between the F-scores from Sensor-seq and the fold enrichment scores from the clonal flow cytometry data (Spearman’s ρ = 0.78) (Supplementary Fig. 12a). In comparison, the results from the secondary screen had only a moderate relationship to the clonal data (Spearman’s ρ = 0.57) (Supplementary Fig. 12b). The poorer correlation likely reflects the inherent noise in screens based on cell sorting. We also examined the false positive and false negative rates at various F-score and fold enrichment thresholds (Supplementary Fig 12c,d). The F-score and fold enrichment values reflect the activity level of individual variants in the RNAseq and sorting experiments, respectively. At a relatively stringent F-score threshold of 3 and a fold enrichment threshold of 1.5, we observed no false positives but ∼41% false negatives. In contrast, a looser F-score threshold of 1.5 and fold enrichment threshold each at 1.5 resulted in ∼7% false positives and ∼19% false negatives. These results suggest a trade-off between false positive and false negative rates at each threshold. Using the same F-score threshold and testing multiple fold enrichment thresholds, we find an average precision (AP) of 0.94 (Supplementary Fig. 12d). Taken together, the pooled data from Sensor-seq largely summarizes the individual measurements acquired from clonal tests despite these errors.

In summary, Sensor-seq enabled the identification of TtgR variants with unique specificity profiles, including rare hits, from the FuncLib ligand-agnostic library. We identified variants exhibiting strong activity on six of the seven non-native ligands, showcasing varying degrees of specificity toward different ligand groups, and enhanced activity on native ligands. These results also highlight the inherent malleability of some proteins such as TtgR to accommodate a diverse family of ligands. We posit that the malleability is a natural consequence of TtgR’s role as a regulator of a multidrug efflux, which necessitates the ability to detect and respond to different compounds, reflecting an evolutionary pressure to maintain a broad ligand-binding capability.

### Unsupervised learning reveals key residues for ligand specificity

We next sought to determine amino acid sequence preferences for each ligand. To uncover and visualize these sequence determinants in the high dimensional sequence-ligand landscape, we generated a two-dimensional Uniform Manifold Approximation and Projection (UMAP) projection of 17,430 variants using physicochemical embeddings^43^. The amino acid at each of the 11 mutated positions in TtgR is incorporated as a feature using a 19-length dimensionally reduced AAindex which captures diverse amino acid properties such as polarity, hydrophobicity, and alpha/beta propensities^44,45^. Thus, each variant is physicochemically encoded with (11 residues x 19 features) 209 total features^45,46^. We then applied Hierarchical Density-Based Spatial Clustering of Applications with Noise (HDBSCAN) to group variants that are similar^47^. With our chosen hyperparameters we were able to organize 16,882 variants (∼97% of total variants) into 23 clusters containing between 93 and 3916 variants, where each cluster represents sequences with similar physicochemical properties (Fig. 3a, Supplementary Figs. 13 and 14, Supplementary Table 5).

**Figure 3.**
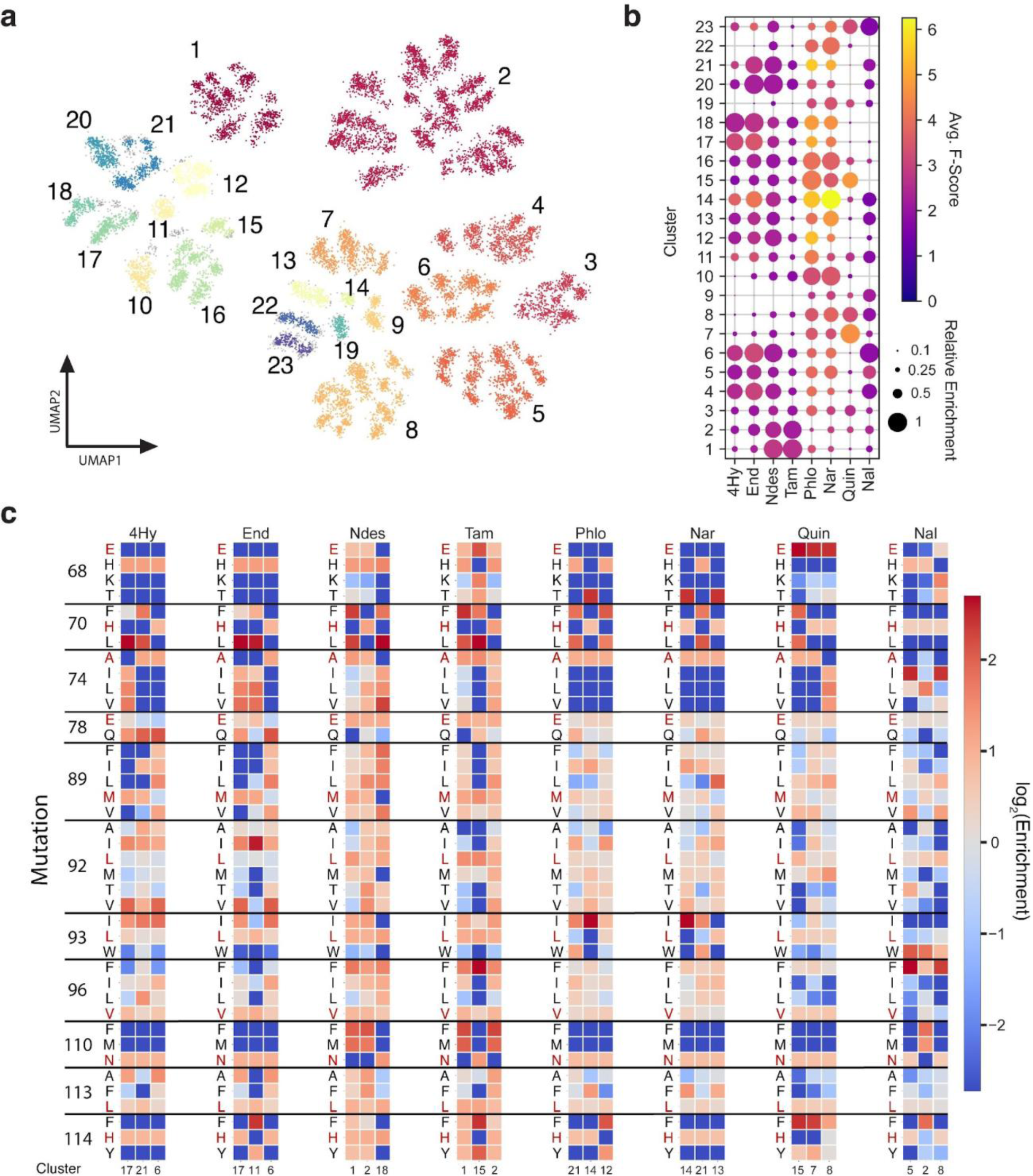
Unsupervised learning reveals amino acid preferences for ligand specificity. ***(a)*** Uniform Manifold Approximation and Projection (UMAP) 2D embedding of 17,430 variants with physicochemical properties of amino acids at each variable position in the TtgR library. Multicolored plot shows 23 clusters identified with Hierarchical Density-Based Spatial Clustering of Applications with Noise (HDBSCAN). Gray points correspond to points the algorithm identified as noise. ***(b)*** Dot plot showing the performance of each of the 23 clusters identified after UMAP-HDBSCAN. The color of each dot represents the average F-score of all variants within the cluster-ligand pair that has a minimum of 1.5 F-score. The size of the dot represents the percentage of variants with >1.5 F-score within each cluster normalized to the highest percentage for that ligand. ***(c)*** Heatmaps for the top 3 performing clusters of each ligand showing log_2_Enrichment of each possible amino acid at the variable positions of TtgR. Clusters are shown from rank 1 to 3 going from left to right. Enrichment was calculated by obtaining the F-score-weighted frequency of amino acids in the cluster using variants with a minimum of 1.5 F-score and normalizing to the DNA count-weighted frequencies of the initial library. Red letters denote the wild-type residues. See Methods for in-depth description of analysis.

To identify functional regions in sequence space for each ligand, we overlaid the F-scores obtained from Sensor-Seq for each ligand on the UMAP projection (Supplementary Fig. 15). An examination of the UMAP with overlaid F-scores revealed that functional sequences for each ligand are distributed across multiple clusters. This implies the potential existence of degenerate solutions, wherein several groups of sequences can bind to the same ligand. A notable feature is the high local ruggedness, where adjacent points within a cluster exhibit gains or losses of function, suggesting that small physicochemical changes in closely related protein sequences are sufficient to alter function dramatically (Supplementary Fig. 15).

To assess the distribution of ligand-responsive sequences across clusters quantitatively, we analyzed the percentage of “hits” and the average F-score of “hits” within a cluster (Fig. 3b, Supplementary Fig. 16, Supplementary Tables 9-10). Here, a “hit” is defined as any variant with an F-score of 1.5 or higher (Fig. 3B, Supplementary Fig. 16). The choice of 1.5 as the threshold for activity represents the approximate value where incremental increases in the threshold do not significantly change the number of hits (Supplementary Fig. 17a,b). Using these metrics, we first compared the performance of each cluster for each ligand to gain a global view of the sequence-function landscape. Then we delved deeper to elucidate sequence determinants of TtgR driving a gain-of-function response to each ligand. This involved examining the enrichment or depletion residues among highly active variants within the top three clusters for each ligand. To ensure that the sequences within the top clusters are a good representation of sequences that would be highly active on a specific ligand, we ranked the clusters based on the average F-score of hits (variants with F-score ≥1.5) and imposed a minimum of 15 hits per cluster.

The hydroxylated tamoxifen derivatives, 4-hydroxytamoxifen, and endoxifen, share similar top-performing clusters (Fig. 3b). As expected, N-desmethyl tamoxifen and tamoxifen share high-performing clusters mostly distinct from the top-performing clusters for 4-hydroxytamoxifen and endoxifen, suggesting, in general, distinct groups of sequences respond to both ligand pairs (Fig 3b). E68H mutation is strongly enriched and L93W is strongly depleted among variants active on all tamoxifen derivatives (Fig. 3c). Variants active on 4-hydroxytamoxifen and endoxifen showed preference for histidine at position 68, glutamine at position 78, and the wildtype residue, asparagine, at position 110 (Fig. 3c). In contrast, the variants active on N-desmethyltamoxifen and tamoxifen were more permissive of different substitutions at position 68, favored glutamate at position 78, and had either a phenylalanine or methionine in top clusters at position 110 (Fig. 3c). These three residues, 68, 78 and 110, may impart specificity for 4-hydroxytamoxifen and endoxifen through hydrogen bonding with the 4-hydroxyl group of the ligand. Specificity could be engineered for 4-hydroxytamoxifen and endoxifen if positions 78 and 110 contain glutamine and asparagine, respectively (Fig. 3c). Notably, Clusters 20 and 21 contain a relatively large portion (12-18%) of sequences that respond to N-desmethyltamoxifen and endoxifen, albeit weakly (lower average F-score than highly active clusters) (Fig 3b), indicating the presence of variants within these clusters that are capable of accommodating both methylated and hydroxylated forms of tamoxifen but not the tertiary amine group found in tamoxifen and 4-hydroxytamoxifen.

Quinine and naltrexone have unique cluster profiles due to their distinct structures (Fig. 3b). Thus, specific protein sequence characteristics govern specificity for each of these ligands. For quinine, we observed enrichment of wildtype residues E68 and N110 and either phenylalanine or tyrosine at position 114, which may stabilize quinine with a pi-stacking interaction. Positions 78, 89, and 92 are tolerant to most substitutions except threonine at position 92 (Fig. 3c). At position 96, isoleucine and leucine are disfavored, but valine and phenylalanine are permissible suggesting that size of the amino acid may not be the determining factor (Fig. 3c). For naltrexone, the key specificity determining mutations are H70, W93, and F96 (Fig. 3c). Interestingly, tryptophan at position 93 is strongly favored for naltrexone, but strongly disfavored for quinine (Fig. 3c).

Naringenin and phloretin-responsive sequences are distributed throughout the 23 clusters suggesting that the response to these native ligands is often robust to mutation (Fig. 3b). Both shared similar mutation profiles with positions 78, 89, 92, 96, and 113 tolerant to several substitutions (Fig. 3c). We observed co-occurring mutations at positions 68 and 70 for both ligands, where H68 is accompanied by either F70 or L70 and T68 is paired with H70 (Fig. 3c). N110 and H114 are strongly preferred for both native ligands (Fig. 3B). Increased activity on the native ligands may be mediated by interplay between these 4 key positions, 68, 70, 110, and 114. Cluster 12, a top cluster for phloretin but not for naringenin, contains sequences with either an H114F or H114Y. We also observed a markedly lower hit percentage for cluster 12 with naringenin (20%) than with phloretin (32%) (Supplementary Table 9). Thus, variants with a substitution at position 114 to an aromatic residue may have shifted activity towards phloretin than naringenin. Indeed, variants 1H8 and 1F6 from our clonal flow cytometry assessment (Fig. 2d) had specificity for phloretin but not naringenin and carried these aromatic substitutions at position 114.

In summary, these results highlight the power of Sensor-seq to provide a holistic view of the mutational adaptability of TtgR to accommodate diverse ligands, and the key determinants of specificity for each ligand. By revealing the distribution of ligand-responsive sequences across clusters and elucidating sequence determinants driving gain-of-function responses, Sensor-seq offers valuable insights into the underlying landscape of protein-ligand interactions. We anticipate that this information-rich dataset will serve as a valuable resource for benchmarking machine learning algorithms aimed at designing and understanding protein-ligand interactions.

### Crystal structure reveals protein-stabilizing motifs for a non-native ligand interaction

To gain structural insights into TtgR’s ligand specificity, we obtained a high-resolution crystal structure of a quadruple variant (A74L, L93W, N110M, and H114F) bound to non-native ligand naltrexone at 2.05Å resolution (Supplementary Table 11). This structure revealed that TtgR’s ability to interact with various ligands is likely due to an unusually large ligand-binding pocket (1500 Å^3^) for a ∼200 residue protein^30^. The substantial pocket volume enables TtgR to bind ligands in various orientations and with different shapes and features, broadening the potential ligands TtgR can bind. For example, in a previous study we showed that TtgR’s binding pocket is malleable enough to bind to resveratrol, in either vertical or horizontal orientations, with distinct sequences^33^. Ligand specificity is achieved by selecting residue rotamer states within the binding pocket that enhance the tightness of ligand fit primarily through van der Waals interactions. This capability is prominently evident in the naltrexone-bound structure. As a charge-neutral molecule, naltrexone does not require complementary charged residues. All four substitutions, A74L, L93W, N110M, and H114F, are small-to-large changes that facilitate better packing of naltrexone within the binding pocket (Fig. 4a). This jigsaw puzzle fit customizes the pocket for naltrexone and excludes other ligands, as seen by the high specificity of this mutant (Fig. 2d, 3A7). For instance, the mutations in the crystal structure appear to be participating in a total of three Methionine-Aromatic (Met-Aro) interactions, a ubiquitous, protein-stabilizing motif, involving the sulfur atom of methionine and the aromatic ring of a partner residue at a molecular distance of ∼4-6 Å, that can play crucial roles in high-affinity ligand binding^48,49^. L93W forms two 5.7-6.2Å Met-Aro interactions with M167 and M89 (Fig. 4b) in the upper section of the binding pocket. N110M and H114F forms a 5.4Å Met-Aro interaction at the lower section of the binding pocket (Fig. 4b).

**Figure 4.**
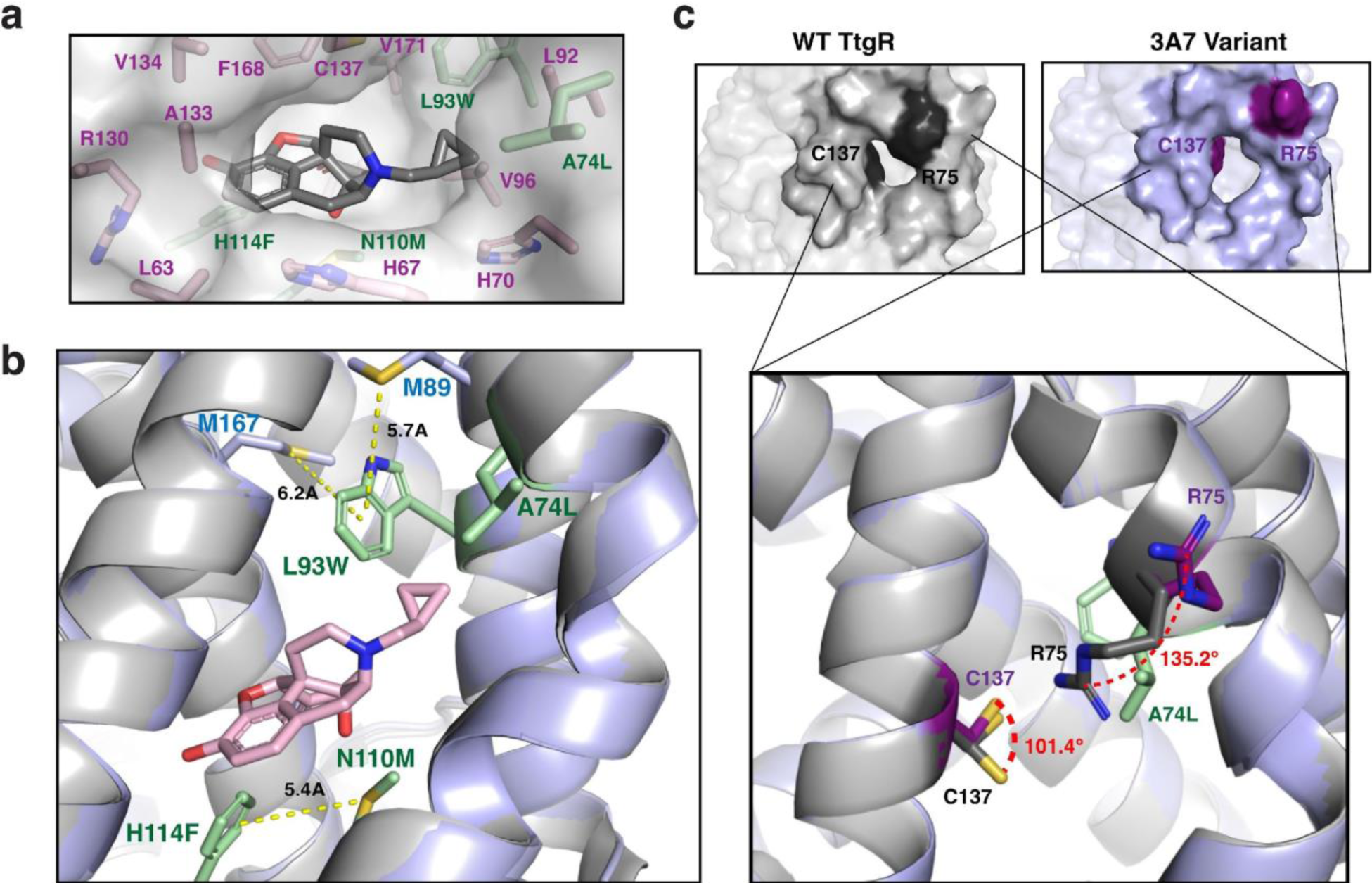
Crystal structure for naltrexone-bound Variant 3A7. ***(a)*** Binding pocket of Variant 3A7 with naltrexone. Gray molecule is naltrexone. Green side chains indicate mutant positions. Pink side chains indicate positions within 4Å of the naltrexone molecule. ***(b)*** Three instances of Met-Aro interactions within the binding pocket, mediated by the mutant residues. WT TtgR (PDB: 7K1C) is shown in gray and 3A7 is shown in light purple. Green side chains indicate mutant positions. Naltrexone is shown in pink. Molecular distances for each interaction are indicated. ***(c)*** Ligand entry port of WT TtgR (gray) and Variant 3A7 (light purple). Darkened surface residues denote positions of C137 and R75. Dotted red lines indicate rotation angles of C137 and R75 in WT (dark gray) TtgR and Variant 3A7 (dark purple). Green side chains indicate mutant positions.

We also observed a larger portal for ligand entry into the binding pocket (Fig. 4c). The entry appears to be widened through the rotation of two key residues, C137 and R75 (Fig. 4c). In the quadruple mutant, the sulfur atom of C137 is rotated maximally ∼101° around the β-carbon from the wildtype rotamer position and towards the entry port, thereby increasing access to the binding pocket^30^. A similar effect can be observed with the R75 guanidino group rotated ∼135° around the β-carbon from the wildtype rotamer position and away from the entry port (Fig. 4c). This rotation can be explained by the additional steric bulk caused by the neighboring A74L substitution (Fig. 4c). The open conformation in 3A7 should allow access of differently-sized molecules to enter the ligand-binding pocket. In summary, the stabilizing energy of the jigsaw-fit ligand-aTF interaction and the larger entry portal likely facilitates the accommodation of naltrexone into the designed TtgR.

### Design of cell-free biosensors for naltrexone and quinine

To demonstrate the practical utility of the designed biosensors, we sought to develop cell-free expression systems (CFE) for simple, economical analyte detection^50,51^. We first tested a biosensor for naltrexone given the compelling need for low-cost point-of-care approaches for detecting opioid use in rural communities without easy access to healthcare^52^. Naltrexone works by binding to and blocking the effects of mu-opioid receptors and is used to treat substance use disorders^53^. As an opioid analog, naltrexone can serve as a stand-in substitute for opioids such as heroin or morphine. Two plasmids – one encoding the naltrexone-responsive 3A7 aTF (sensor plasmid) and one encoding GFP regulated by a TtgR promoter (reporter plasmid) – were added to a cell-free reaction containing processed *E. coli* extract together with the additional biological cofactors required for transcription and translation *in vitro*^54^ (Fig. 5a). We first validated the ability of the sensor to repress expression of the fluorescent reporter by testing a range of sensor plasmid concentrations (0-10nM) in the absence of analyte and quantifying fluorescence over 18 hours (Fig. 5b). We observed dose-dependent repression of the GFP signal, with fluorescence being distinguishable across sensor plasmid concentrations within 2 hours and reaching steady-state levels at ∼10-12 hours (Fig. 5b). The minimum fluorescence level was reached at 5nM sensor plasmid (Fig. 5b). Having validated the 3A7 sensor as a functional repressor in a CFE, we next tested whether varying concentrations of naltrexone (0-100µM) can be detected by the sensor and abolish repression. Generally, by increasing the amount of naltrexone, we observed higher steady-state levels of fluorescence, with the largest stepwise increase obtained transitioning from 10nM to 100nM (Fig. 5c). At 100µM naltrexone, we observed higher variance of the measurements, suggesting possible inhibition of the cell-free reaction at high concentrations of analyte (Fig. 5c). The fluorescence signal was readily distinguishable between 10nM naltrexone and higher concentrations within 2 hours, but took longer at lower concentrations (Fig. 5c).

**Figure 5.**
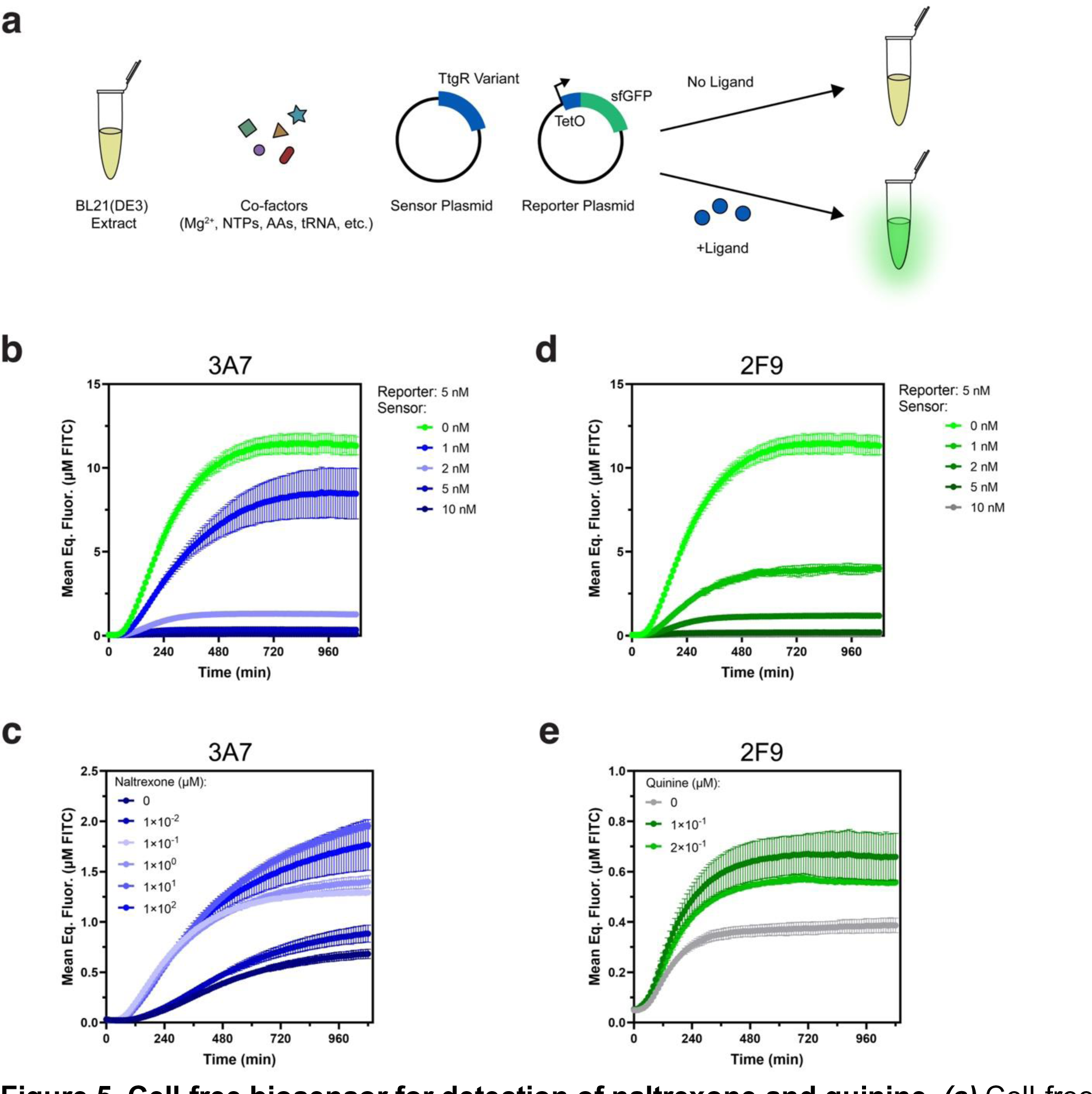
Cell-free biosensor for detection of naltrexone and quinine. ***(a)*** Cell-free expression system consisting of a sensor plasmid and a sfGFP reporter plasmid whose expression is mediated by ligand binding by the sensor. ***(b)*** The plasmid encoding 3A7 TtgR (sensor plasmid) was titrated against 5 nM of the plasmid encoding sfGFP regulated by 3A7 TtgR (reporter plasmid) in cell-free gene expression reactions. ***(c)*** 6 nM of the 3A7 sensor plasmid and 2 nM reporter plasmid were added to cell-free gene expression reactions and incubated with naltrexone at log-fold increments. ***(d)*** 2F9 TtgR sensor plasmid was titrated against 5 nM reporter plasmid in cell-free gene expression reactions. ***(e)*** 4 nM 2F9 TtgR sensor plasmid and 2 nM reporter plasmid were added to cell-free gene expression reactions and incubated with indicated concentrations of quinine. Reported are the sfGFP concentrations normalized to a FITC standard over 18-hours experiment at 30 °C, continuously monitored every 10 minutes. Points indicate individual replicates (N = 3).

Next, we aimed to create a cell-free sensor for quinine, another non-native ligand. Quinine has been used since the 17^th^ century as an antimalarial drug and, despite its well-documented adverse side effects, is still in use today, primarily in underdeveloped countries with resource-limited healthcare systems or where safer modern alternatives are not available^40^. In such communities, significant levels of quinine are likely to be detected in the wastewater, which is often recycled for irrigation or freshwater sustainability and can carry unforeseen environmental and health ramifications^55^. Thus, an affordable biosensor for monitoring quinine metabolism and detection in water supplies may be of interest. We tested the 2F9 variant, one of the strongest quinine responders from our tertiary validation. We examined the time-dependent changes in fluorescence after adding a range of sensor plasmid (0-10nM) in a similar experimental set-up with the naltrexone sensor. Like with the 3A7 sensor, we observed a dose-dependent repression that was evident within 2 hours, and we found that 5nM of sensor plasmid was sufficient for minimal fluorescence (Fig. 5d). Having validated that the 2F9 sensor can repress reporter expression, we next tested if expression can be restored with the addition of quinine. We tested 3 concentrations of quinine (0uM, 100nM, and 200nM) for 18 hours. We found that at 100nM, the initial fluorescence difference was distinguishable in two hours and approximately doubled at steady-state, which took ∼6-8 hours to reach (Fig 5e). We also observed a decrease in overall fluorescence when the quinine concentration was increased to 200nM (Fig. 5e). Quinine has been reported to be a weak intercalator of DNA, thus high concentrations may have inhibitory effects on expression of the GFP reporter^56^. These data are a promising step toward developing cell-free assays for naltrexone and quinine detection. Future steps could optimize the cell-free biosensors in matrices such as wastewater and biological fluids,

## Discussion

Sensor-seq is a platform technology that can engineer aTFs with novel ligand specificities. By integrating phylogeny-guided sequence diversification to preserve allosteric signaling with an RNA barcoding system, Sensor-seq screens aTF variants through deep sequencing to achieve both sensitivity and scale. When applied to redesign the specificity of a bacterial aTF, TtgR, Sensor-seq yielded variants exhibiting strong activity on six of seven non-native ligands, with distinct specificity profiles toward different ligand groups and enhanced activity and specificity toward native ligands. Statistical analysis of this dataset comprising nearly 160,000 sequence–function data points elucidated sequence determinants driving gain-of-function and provided detailed insights into the underlying landscape of protein-ligand interactions.

The evolvability of TtgR, as indicated by its remarkable adaptability to acquire novel functions, can be understood in the context of its native role as regulator of the TtgABC multi-drug efflux pump operon^30,57^. TtgR’s role in regulating a multi-drug exporter necessitates its ability to detect and respond to different compounds, reflecting an evolutionary pressure to maintain a broad ligand-binding capability. This natural adaptive potential can be advantageous for the evolution of new functions. From a structural standpoint, TtgR’s multi-functionality arises from an unusually large binding pocket volume that allows TtgR to accommodate ligands of various sizes and orientations^30,33^. Ligand specificity is achieved by the selection of residue rotamer states within the binding pocket that enhance the tightness of ligand fit primarily through van der Waals interactions. Other bacterial regulators of multi-drug efflux pumps may emerge as potential scaffolds for future biosensor design studies^58^.

Redesigning the ligand specificity of allosteric proteins is challenging due to the interconnectedness of residues involved in ligand binding and those crucial for allosteric actuation. Sequence diversification must carefully balance both generating the necessary diversity for new ligand recognition as well as accounting for allosteric hotspots required for function^59^. A key factor in this balance is the FuncLib method for library generation.

FuncLib excludes mutations that are unlikely to occur naturally. This important step eliminates mutations that are likely to disrupt allostery and structure but preserves mutations that provide diversity. Prior studies on the TtgR family regulators, such as TetR, showed that a mutation that disrupts allostery can be rescued by a compensatory mutation elsewhere^22^. It is possible that mutations generated using FuncLib preferentially select for combinations that lead to allosterically functional proteins. These combinatorial mutations may allow TtgR to sample new conformational states that may confer new ligand specificities without abrogating native function and may explain the extraordinary diversity of ligands binding to TetR-family proteins^1,60^.

Resources such as PDB or sequence databases have facilitated the development of machine learning tools for the prediction of structure and mutational effects on proteins^61–63^. However, large-scale experimental datasets of designed protein–ligand interactions do not exist. The dataset of ∼160,000 quantitative, sequence–function relationships from this work represents a useful large-scale experimental study of protein–ligand specificities of an allosteric protein. Despite focusing on a single protein scaffold, the dataset’s significance lies in its potential utility for training machine learning models. These models could, in turn, predict TtgR sequences capable of binding to novel ligands absent from the original training set. As additional scaffolds are incorporated in future studies, we envision these datasets evolving into essential benchmarks for refining and evaluating machine learning algorithms dedicated to deciphering and designing protein–ligand interactions.

Sensor-seq is a general platform for discovery and design. Beyond its role in engineering functions, it serves as a useful tool for advancing basic science studies by integrating diverse protein libraries, generated through methodologies like deep mutational scanning, ancestral sequence reconstruction, and metagenomic analysis of protein domains to study sequence–function relationships and evolutionary biochemistry. Moreover, Sensor-seq’s scope extends beyond one-component transcription factors (aTFs) to encompass multi-protein relays involved in transcription, such as two-component systems, chemoreceptors, and quorum sensing. This adaptability positions Sensor-seq as a comprehensive and adaptable resource for both applied engineering and the elucidation of fundamental biological principles.

**Supplementary Figure 1.**
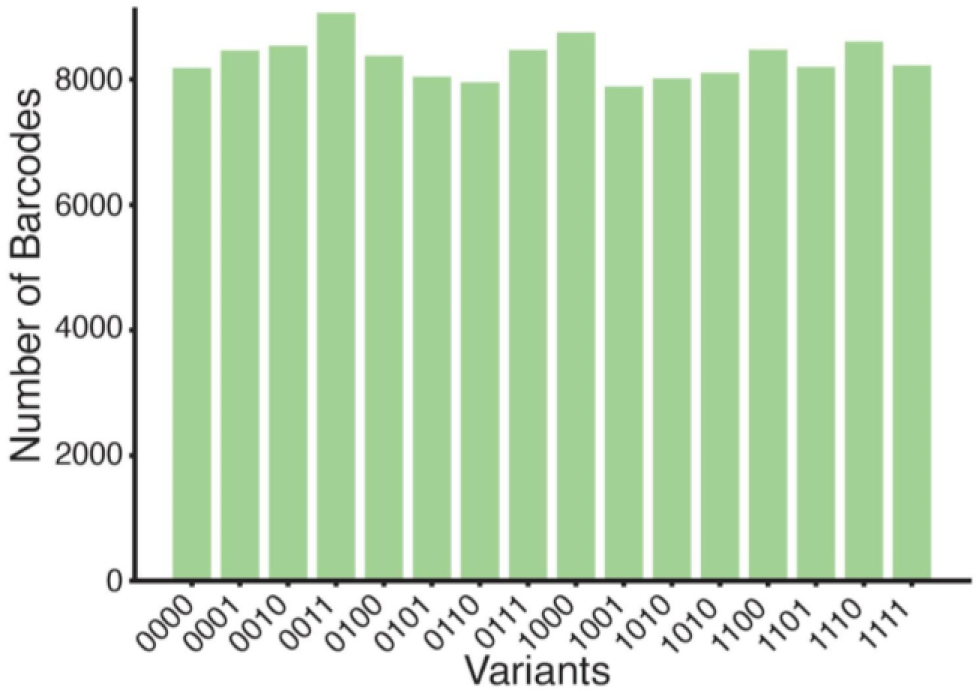
Barcode counts for 16 TtgR variants used in Sensor-seq pilot study with naringenin. The number of barcodes per variant identified using next-generation sequencing. Each variant is identified by a separate binary string.

**Supplementary Figure 2.**
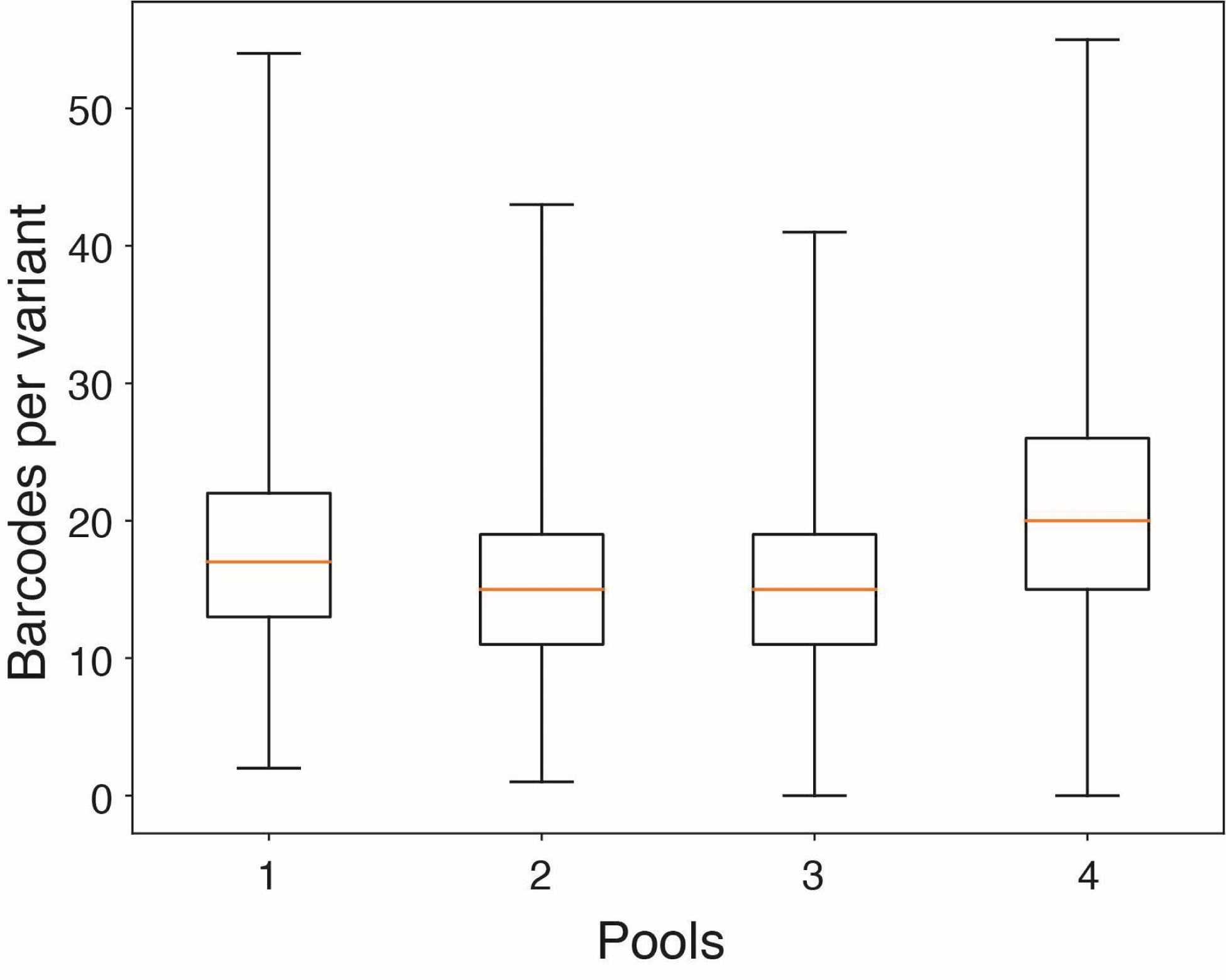
Barcodes per variant of agnostic library from mapping data. The agnostic library was split into 4 pools containing approximately 4,400 variants (see methods). Barcodes and variants were mapped as separate pools. The box plot represents the number of barcodes mapped to each variant in the pools.

**Supplementary Figure 3.**
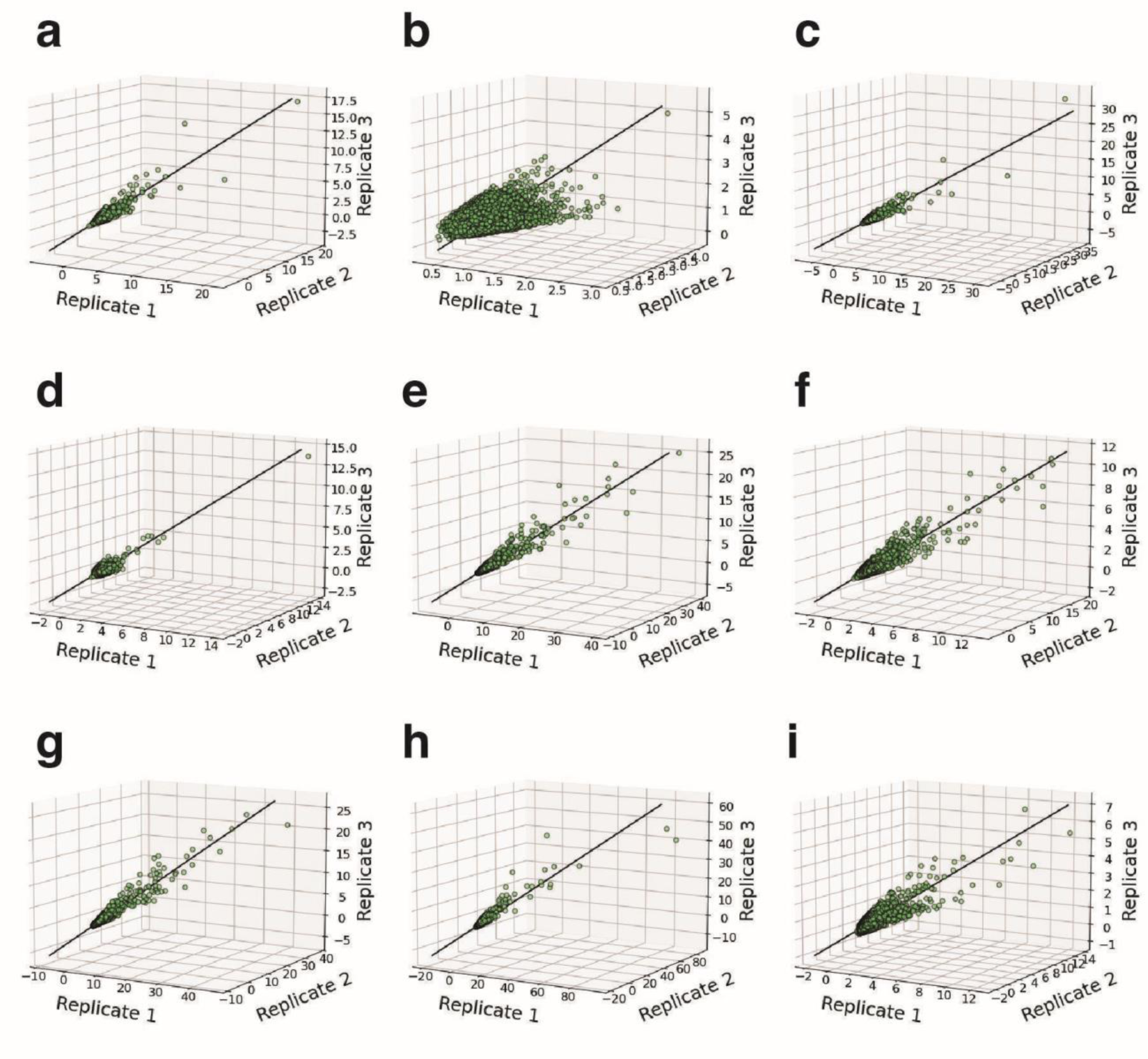
CV filter of RNA-Seq fold enrichment for the agnostic library. Scatter plots of agnostic variants after applying a 30% CV filter for (a) 4-hydroxytamoxifen, (b) ellagic acid, (c) endoxifen, (d) naltrexone, (e) naringenin, (f) N-desmethyltamoxifen, (g) phloretin, (h) quinine, and (i) tamoxifen. The fold enrichment of each variant (green circle) is plotted across the three biological replicates on the X, Y, and Z axes. The best fit line is shown in black and is calculated using a least squares approach.

**Supplementary Figure 4.**
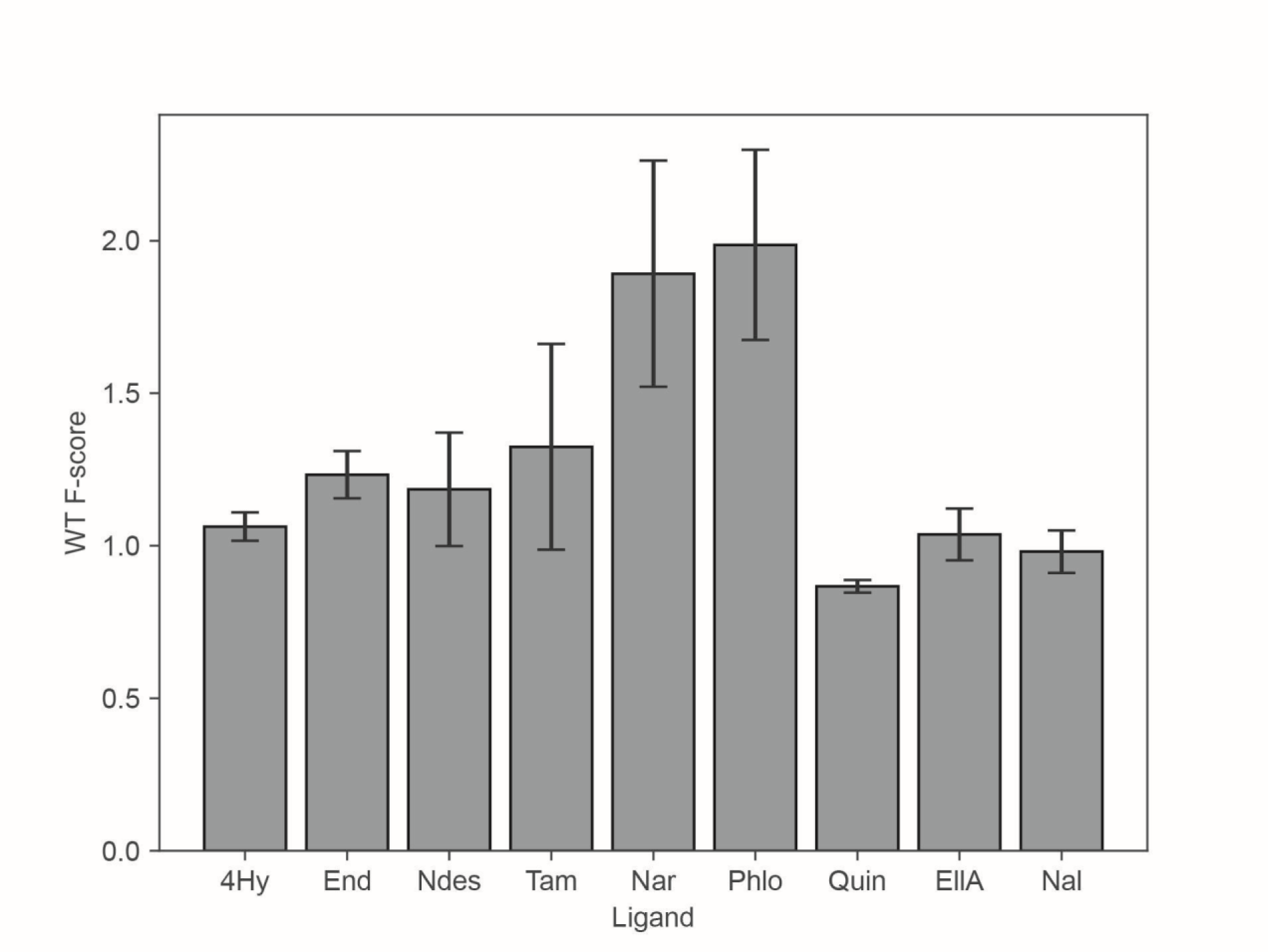
WT TtgR F-scores. F-scores for WT TtgR obtained after Sensor-seq on each ligand. Bars represent an average of three replicates after F-score calculation (see Methods). Error bars represent standard deviation of the three replicates.

**Supplementary Figure 5.**
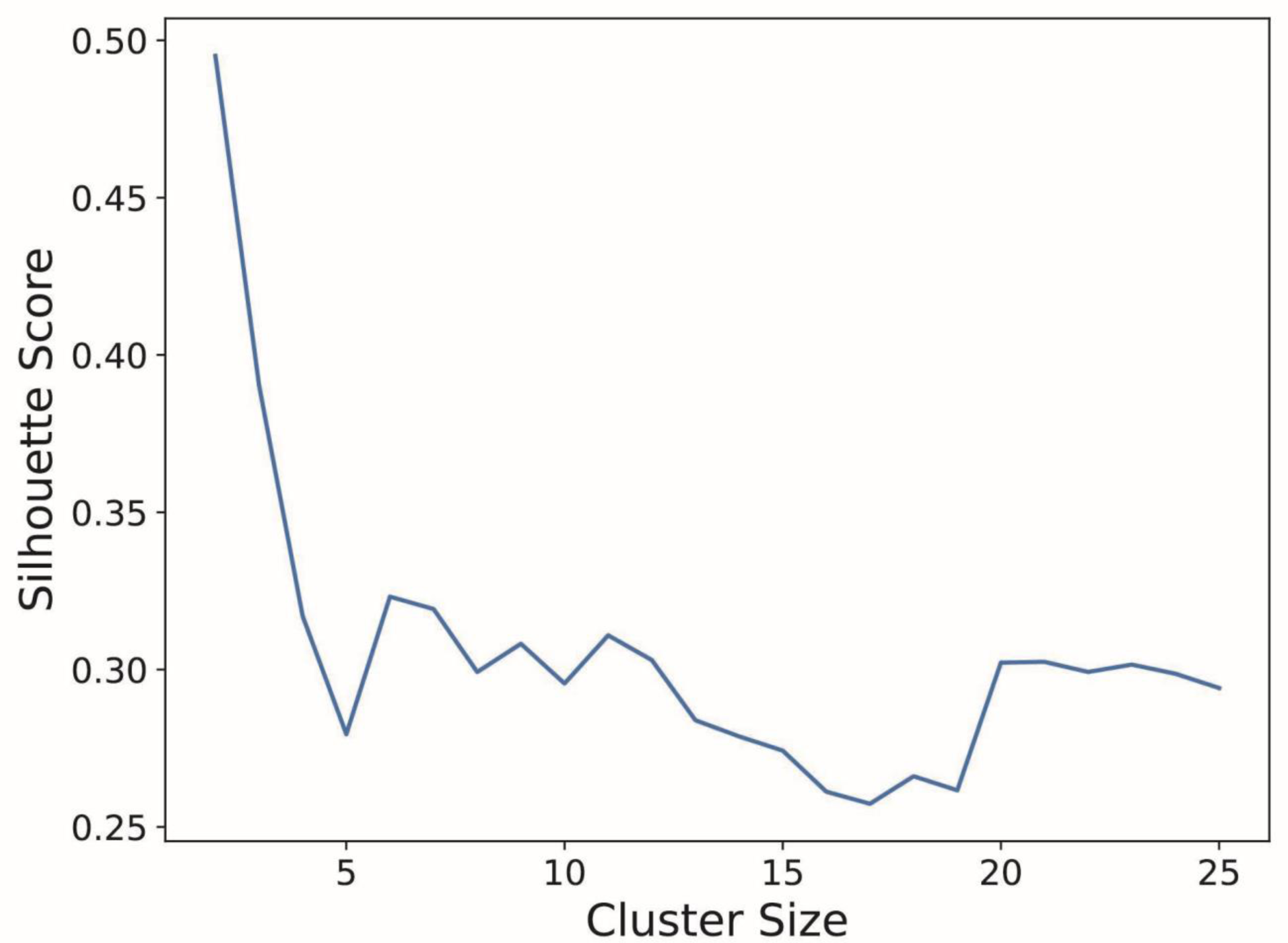
Silhouette score of cluster sizes. The silhouette score is plotted against cluster size for hierarchical clustering with the UPGMA algorithm with a correlation distance metric (see Methods).

**Supplementary Figure 6.**
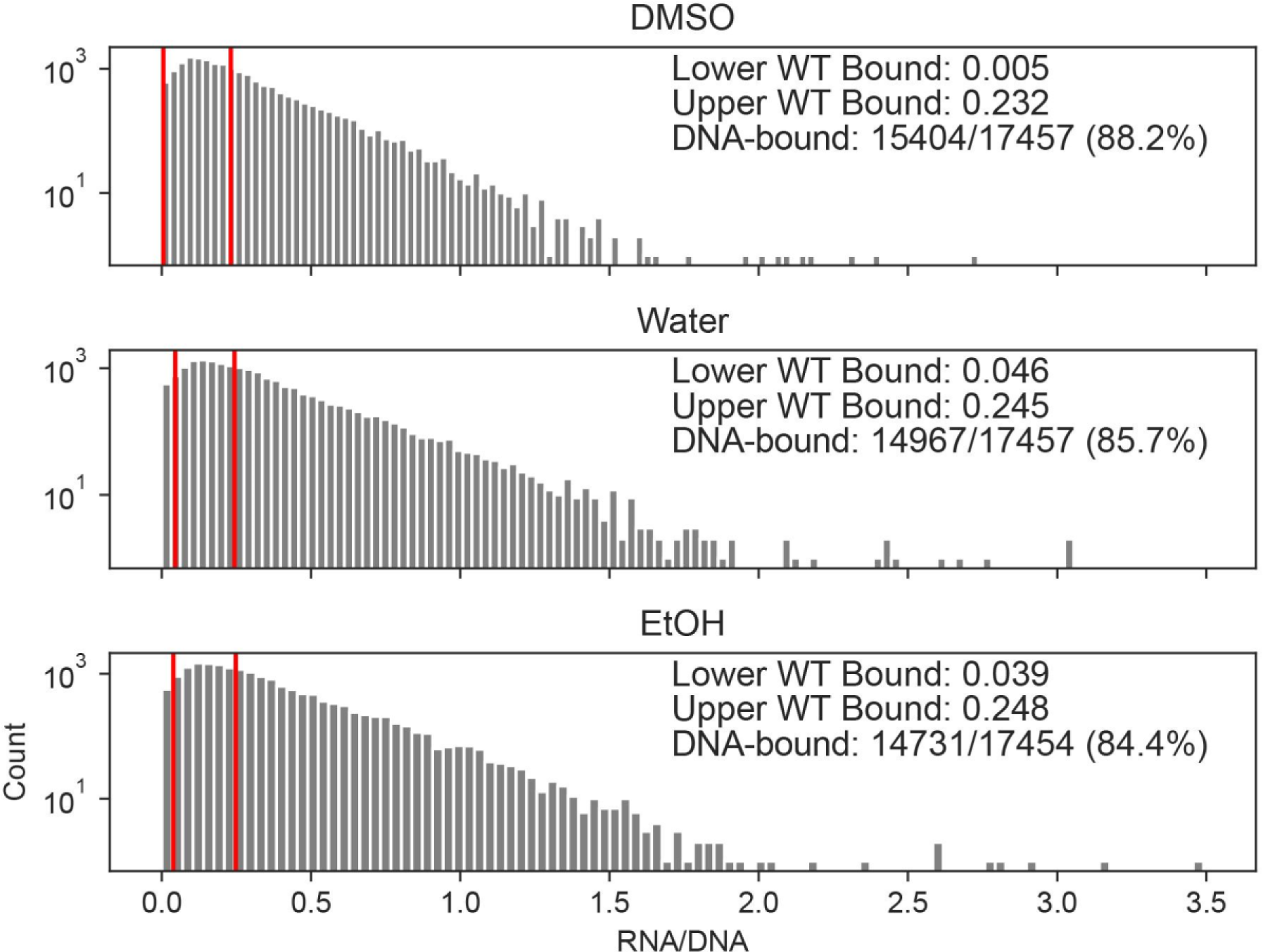
Quantification of DNA-bound TtgR variants (functional repressors). The RNA/DNA count ratio for each variant with each solvent was calculated and the means of three replicates were plotted in a histogram. The 95% confidence interval for WT TtgR ratio under each solvent is denoted by the red lines. We define this interval for WT TtgR as repressor-competent (i.e. able to bind DNA and repress transcription). 95% confidence intervals for each variant were then calculated and those that overlap with the WT intervals were identified as DNA-bound and therefore, functional repressors.

**Supplementary Figure 7.**
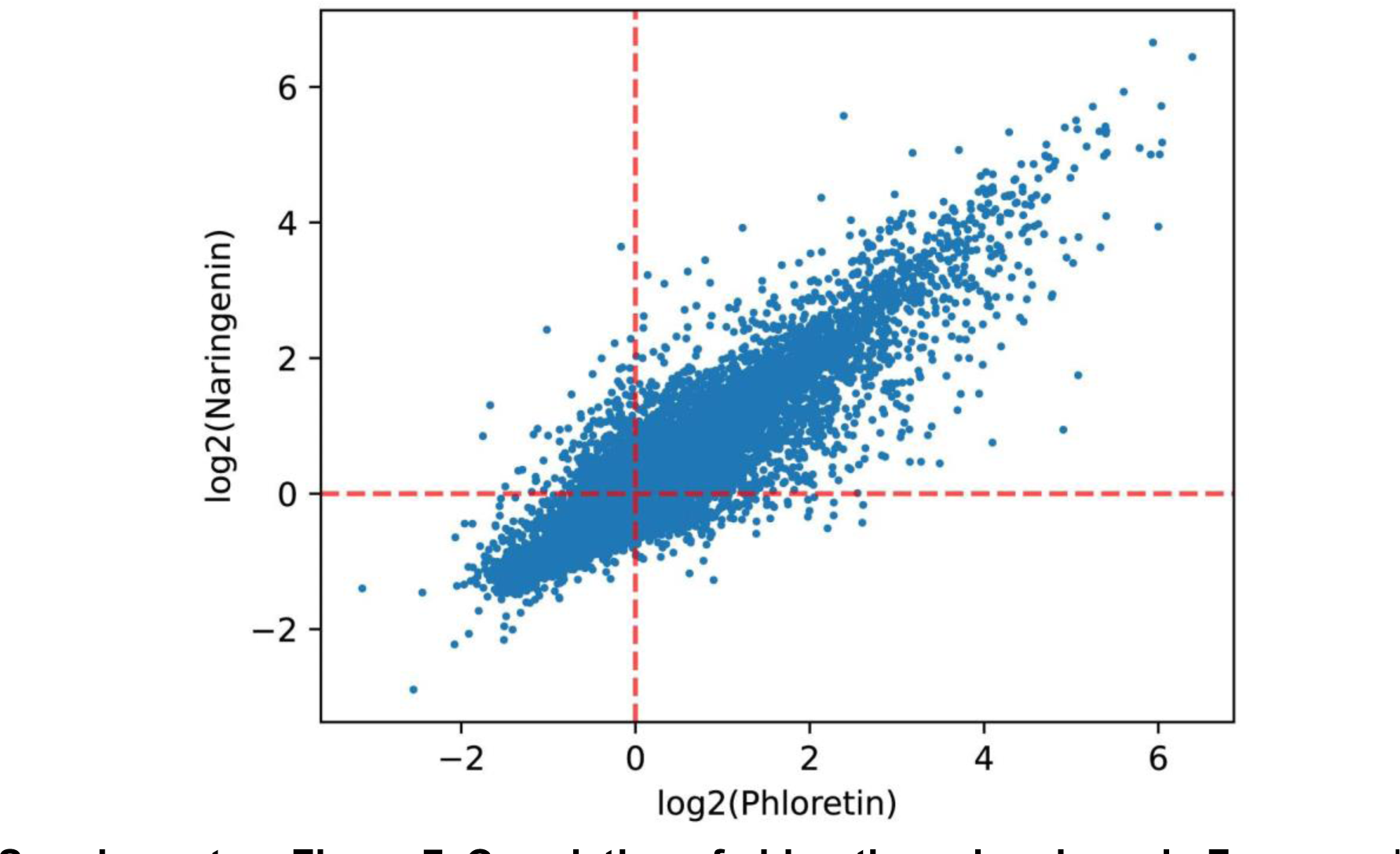
Correlation of phloretin and naringenin F-scores. Log_2_ transformed F-scores for all variants under phloretin and naringenin are plotted. Red dotted line corresponds to F-score of 1 (no activity).

**Supplementary Figure 8.**
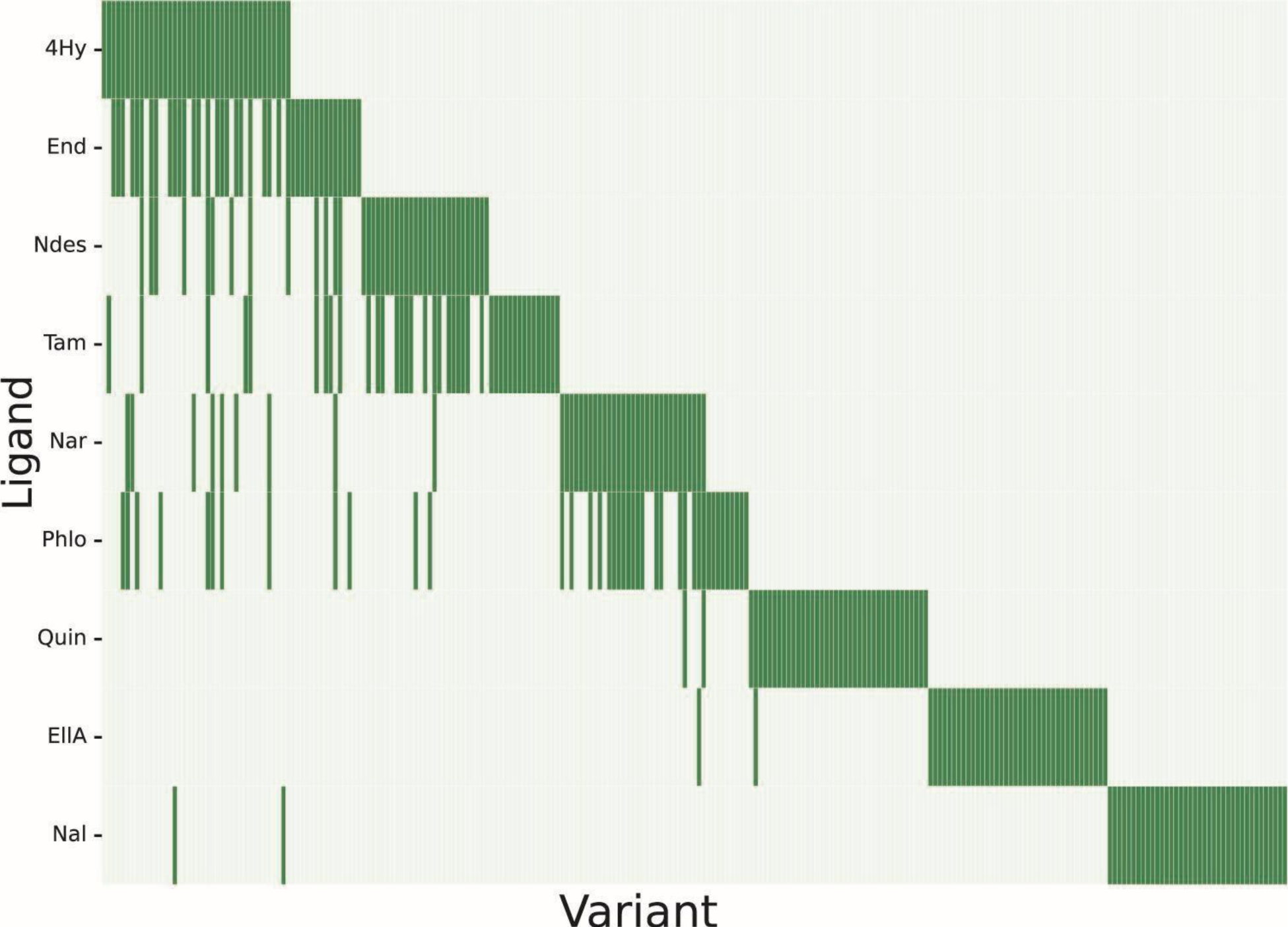
Shared sequences in top performing variants. The top 40 variants for each ligand were selected and listed (x-axis). Variants are marked in green if they are within the top 40 for a particular ligand. There are 251 unique variants in the set of top performers across all ligands.

**Supplementary Figure 9.**
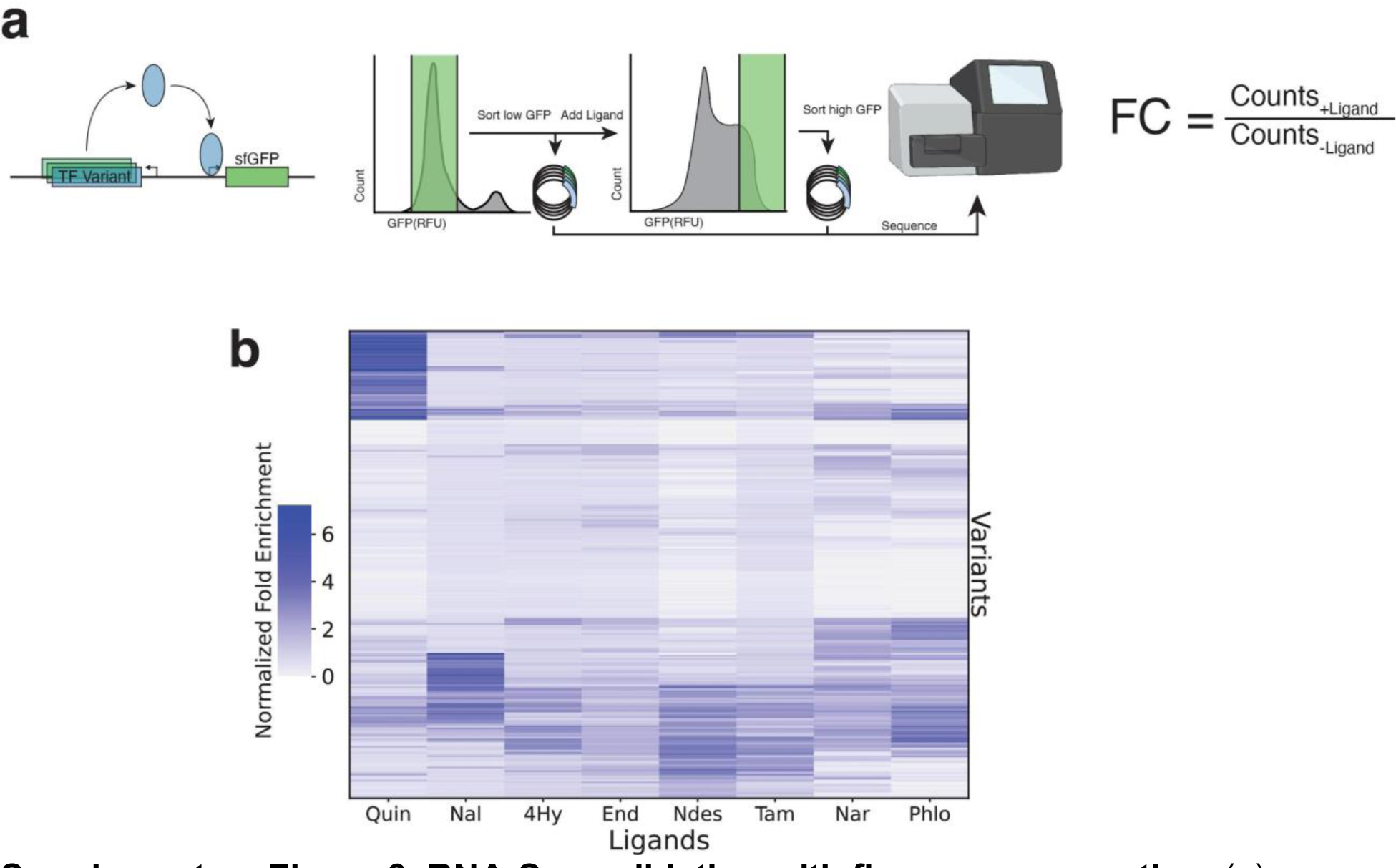
RNA-Seq validation with fluorescence sorting. (a) Fluorescence screening workflow that incorporates a construct with sfGFP. A single repressed sort and an induced sort were sequenced (see methods). Fold change (FC) was calculated as the ratio of percent change in the population with and without ligand. (b) Fold enrichment for each ligand across the 251 best performing variants.

**Supplementary Figure 10.**
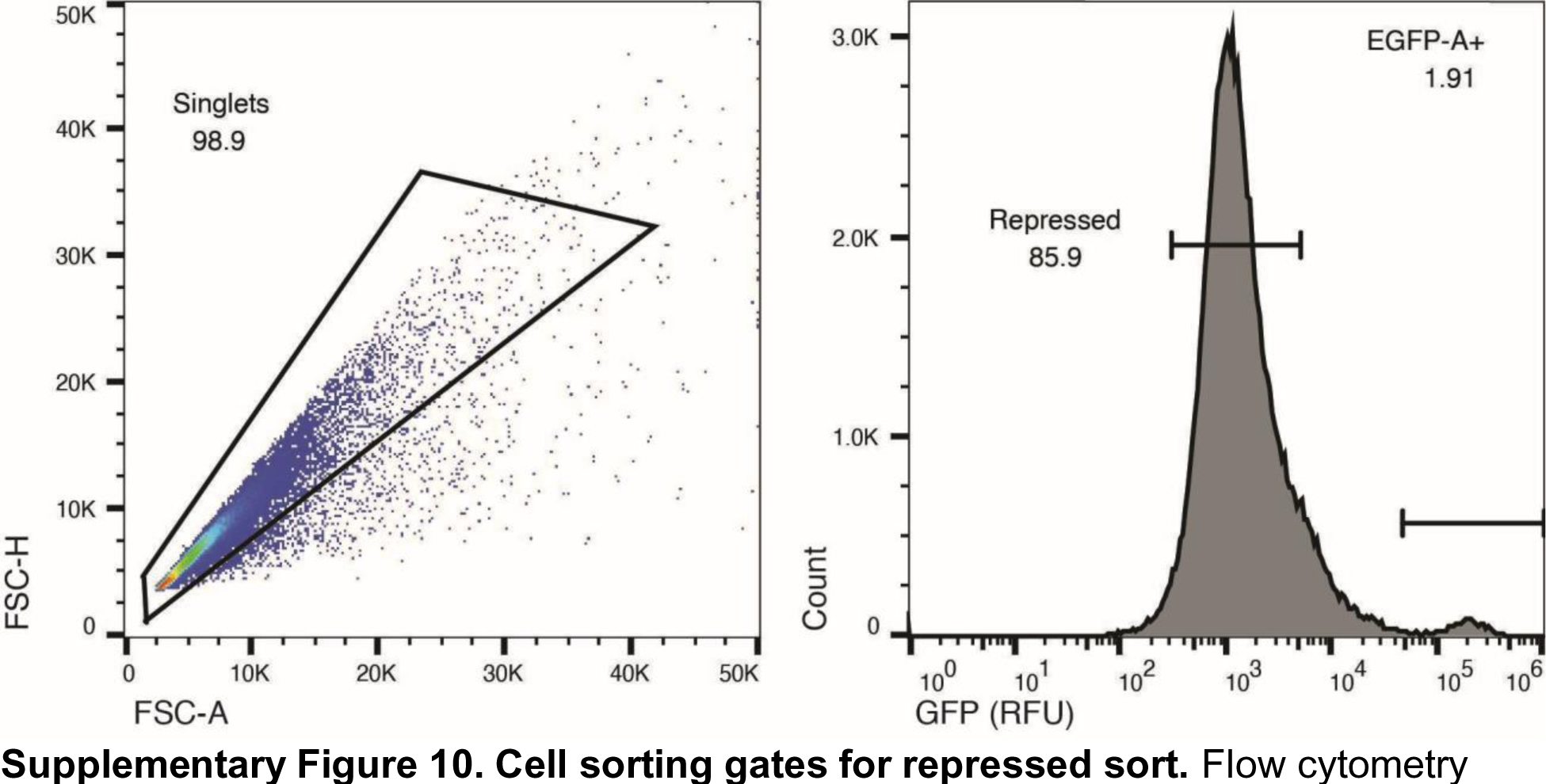
Cell sorting gates for repressed sort. Flow cytometry scatterplot (left) and fluorescence histogram (right) for the library of top variants with no ligand. The scatterplot shows forward scatter area (FSC-A) and forward scatter height (FSC-H). The singlet gate was used to subset the population. Cells falling into the repressed gate in the fluorescence histogram were sorted.

**Supplementary Figure 11.**
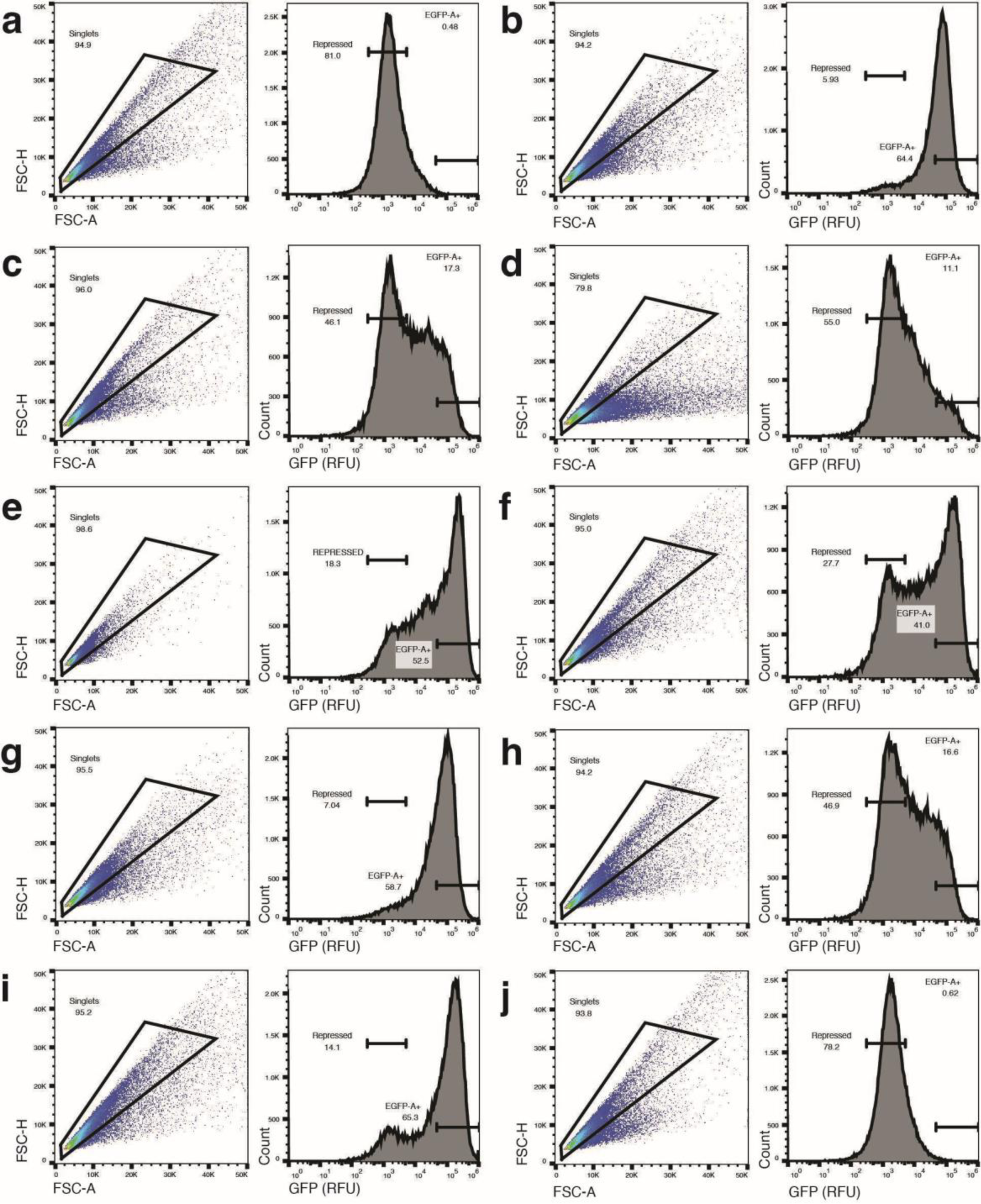
Cell sorting gates for ligand-induced cultures. Flow cytometry scatter plots and histograms for libraries induced with (a) no ligand, (b) naringenin, (c) tamoxifen, (d) naltrexone, (e) quinine, (f) endoxifen, (g) phloretin, (h) 4-hydroxytamoxifen, (i) N-desmethyltamoxifen, (j) ellagic acid. The scatterplot shows forward scatter height versus area; cells falling into the singlet gates were sorted based on the fluorescence histogram (EGFP-A+ gate). The repressed gate indicates the major peak of the no ligand population.

**Supplementary Figure 12.**
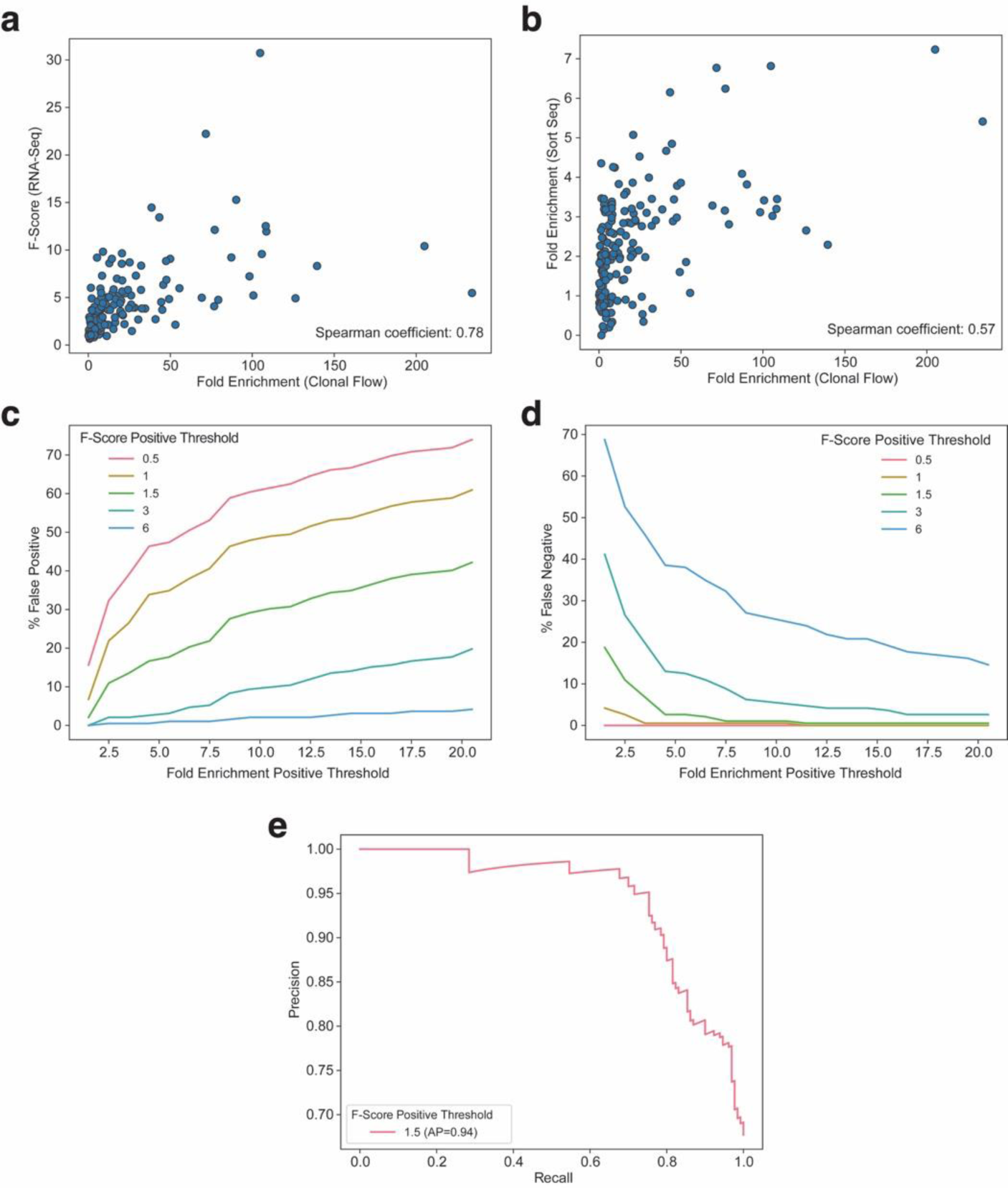
Evaluation of Sensor-seq performance. The fold enrichment scores calculated from the clonal validation of 24 variants across 8 ligands were used as the “ground truth” to assess Sensor-seq performance. (a) Scatter plot depicting the Spearman’s rank correlation coefficient between the clonal flow data and the RNA-seq data. (b) Scatter plot depicting the Spearman’s rank correlation coefficient between the clonal flow data and the secondary screen (sorting and sequencing). (c) Percentage of false positives found within dataset were calculated by varying the positive thresholds for the fold enrichment (clonal flow data) and the F-Score (RNA-seq data). A false positive is defined as any instance where a variant’s F-Score is greater than the F-Score threshold, but its fold enrichment is lower than the fold enrichment threshold. (d) Percentage of false negatives found within dataset were calculated by varying the positive thresholds for the fold enrichment (clonal flow data) and the F-Score (RNA-seq data). A false negative is defined as any instance where a variant’s F-Score is less than the F-Score threshold, but its fold enrichment is greater than the fold enrichment threshold. (e) Precision-recall curve varying the fold enrichment thresholds at a constant F-Score positive threshold of 1.5.

**Supplementary Figure 13.**
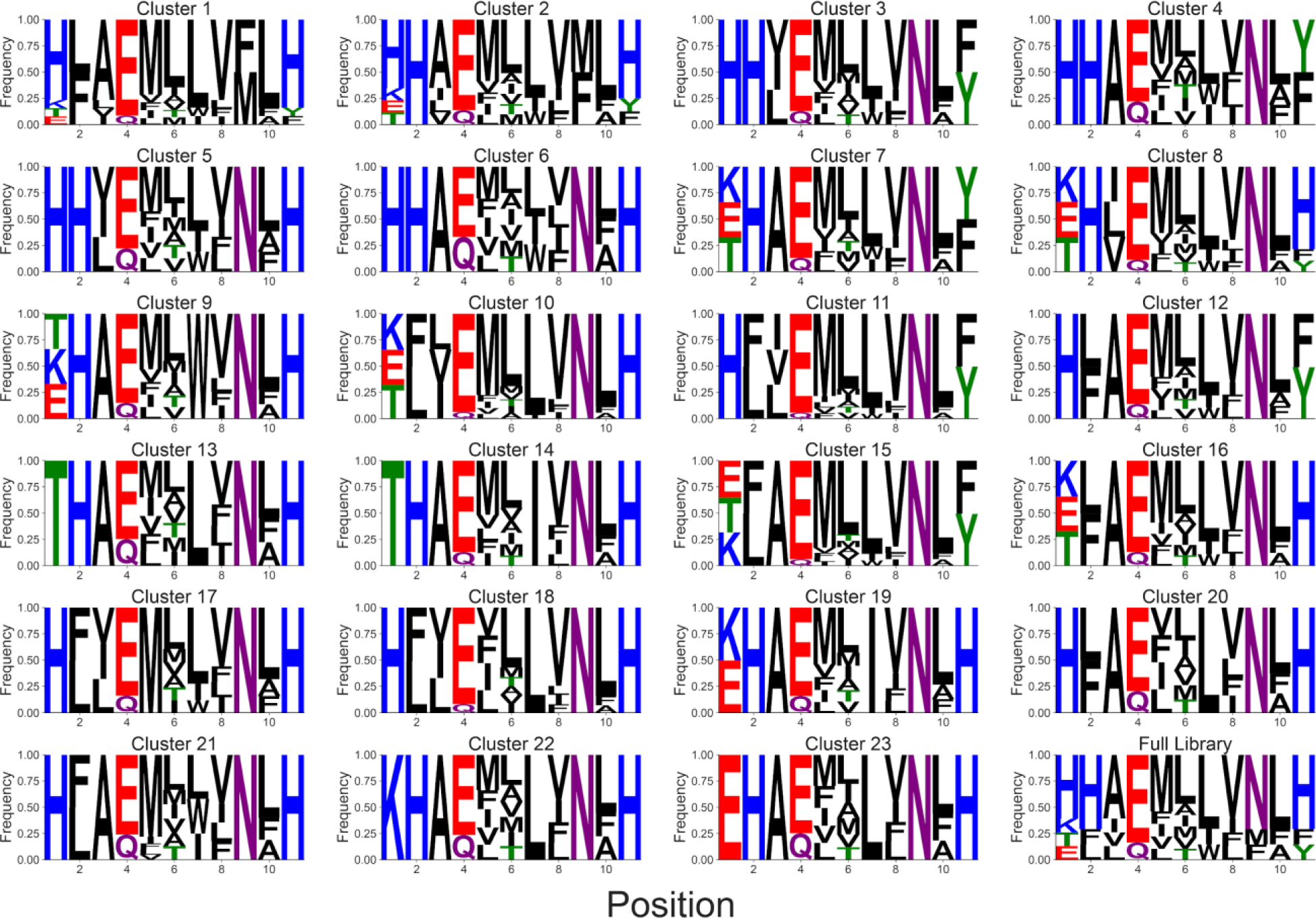
Sequence logos for HDBSCAN clusters. Sequence logos showing frequencies of each amino acid at each position after HDBSCAN of a UMAP embedding with the following paramters: 500 n_neighbors, 0.1 min_dist, 80 min_cluster_size (see Methods). Amino acids are colored based on functional groups. Logos were made using LogoMaker^64^.

**Supplementary Figure 14.**
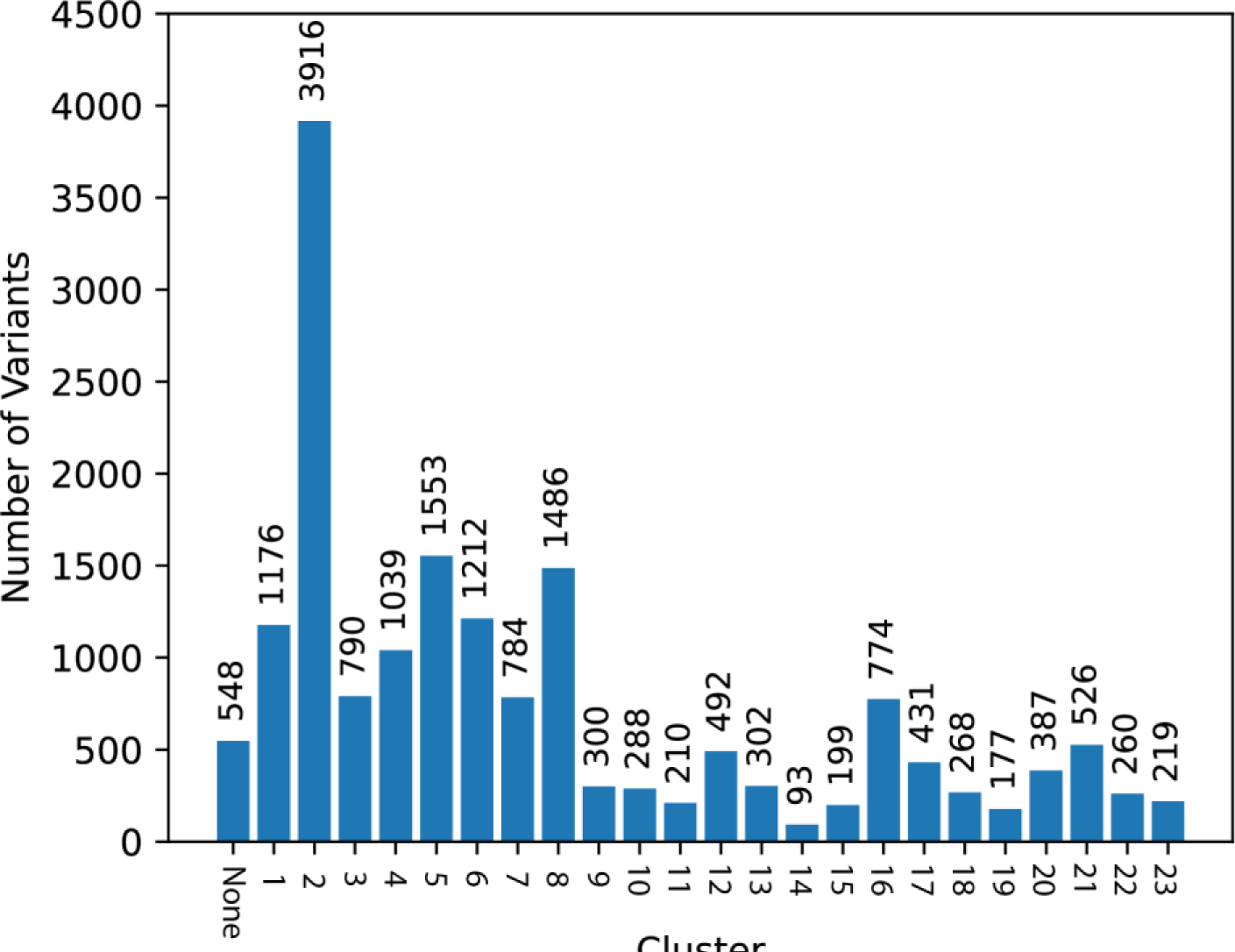
HDBSCAN cluster sizes. Cluster sizes after HDBSCAN of a UMAP embedding with the following parameters: 500 n_neighbors, 0.1 min_dist, 80 min_cluster_size (see methods).

**Supplementary Figure 15.**
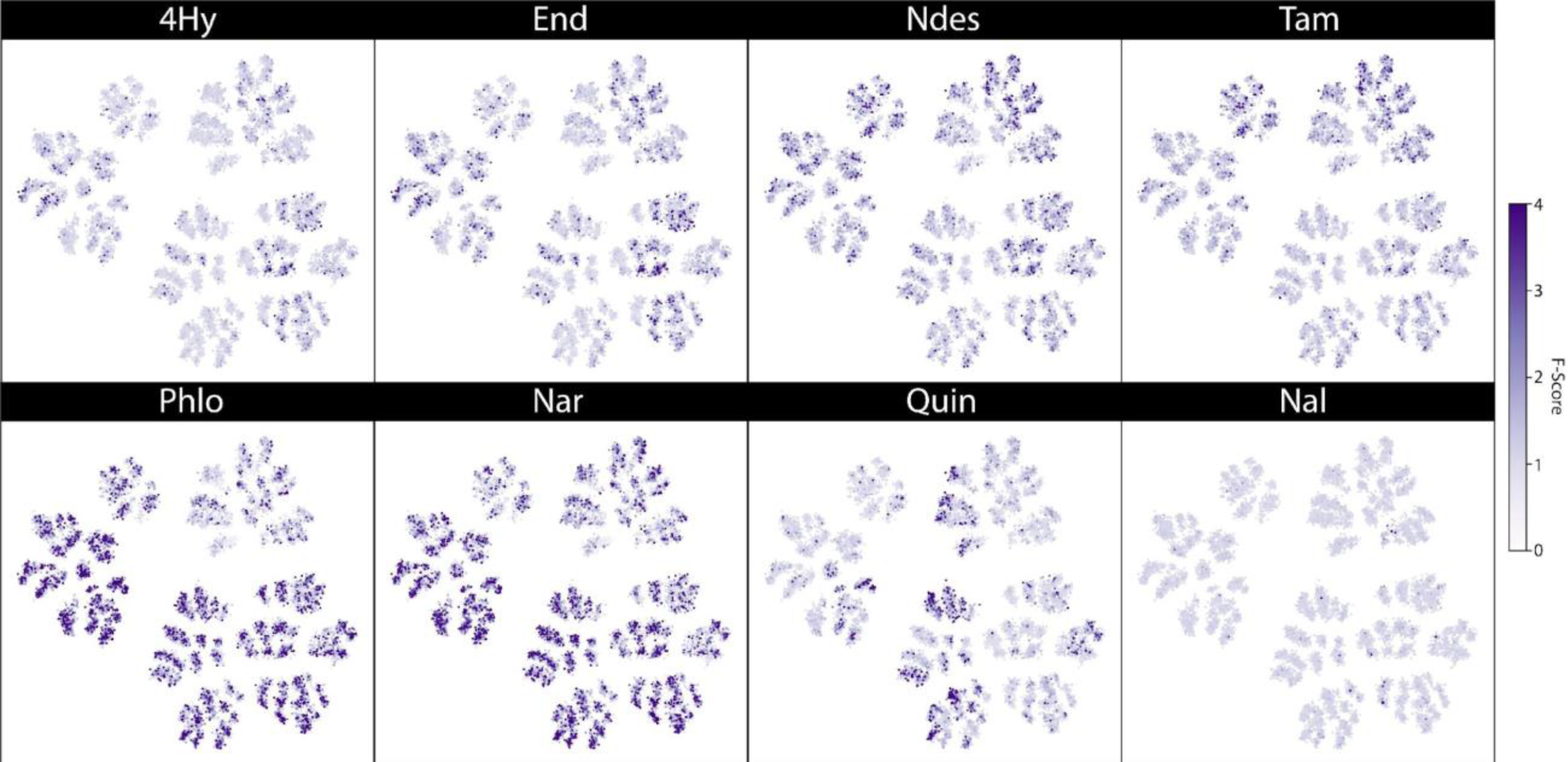
Overlay of F-scores on UMAP projection. UMAP embedding was generated with the following parameters: 500 n_neighbors, 0.1 min_dist, 80 min_cluster_size. The F-scores of each variant under each ligand calculated from Sensor-seq are represented by the shade of each point. An F-score ceiling of 4 was imposed for shading.

**Supplementary Figure 16.**
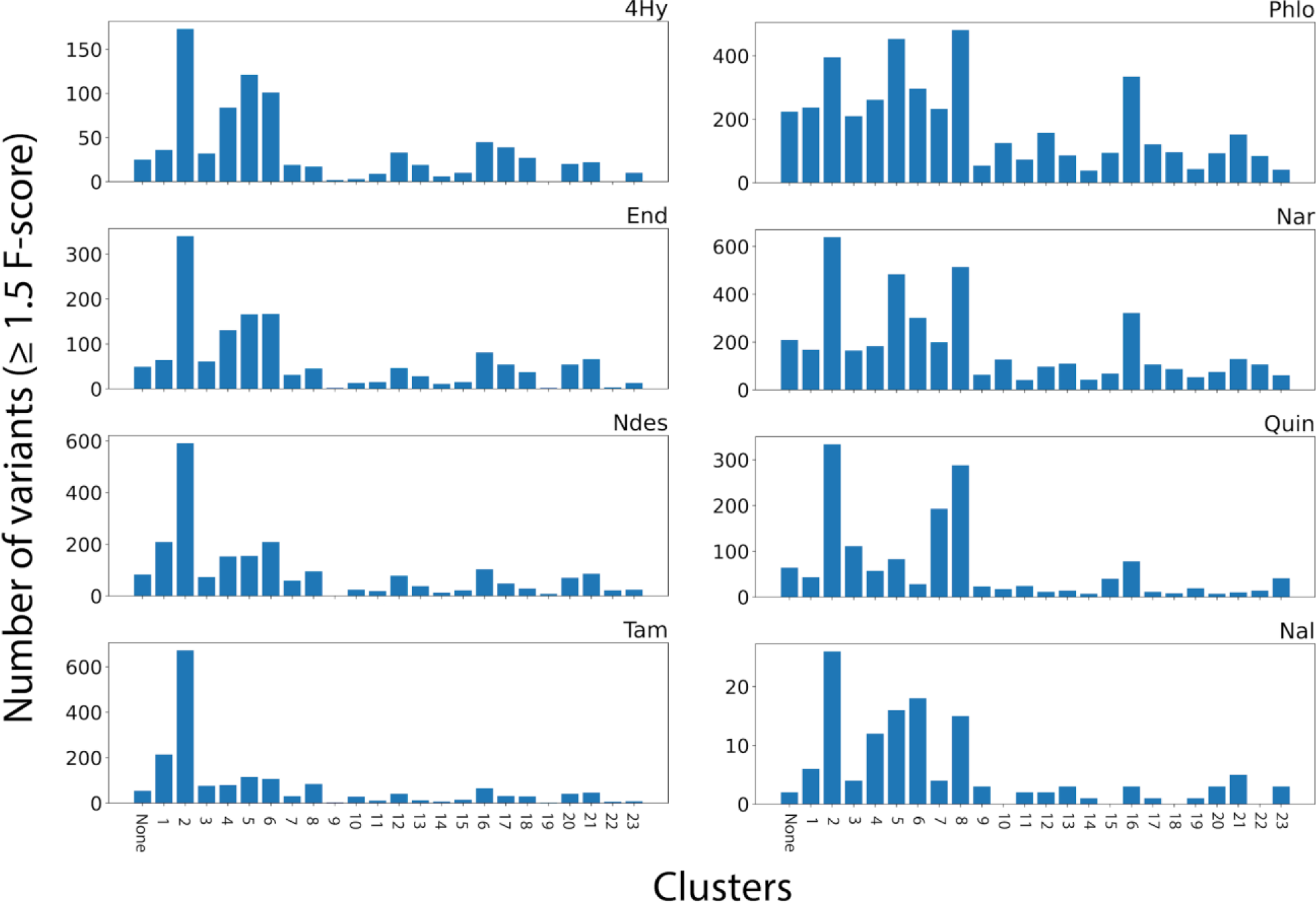
HDBSCAN cluster sizes passing F-score filter. Cluster sizes containing only variants with ≥1.5 F-score after HDBSCAN of a UMAP embedding with the following parameters: 500 n_neighbors, 0.1 min_dist, 80 min_cluster_size (see methods).

**Supplementary Figure 17.**
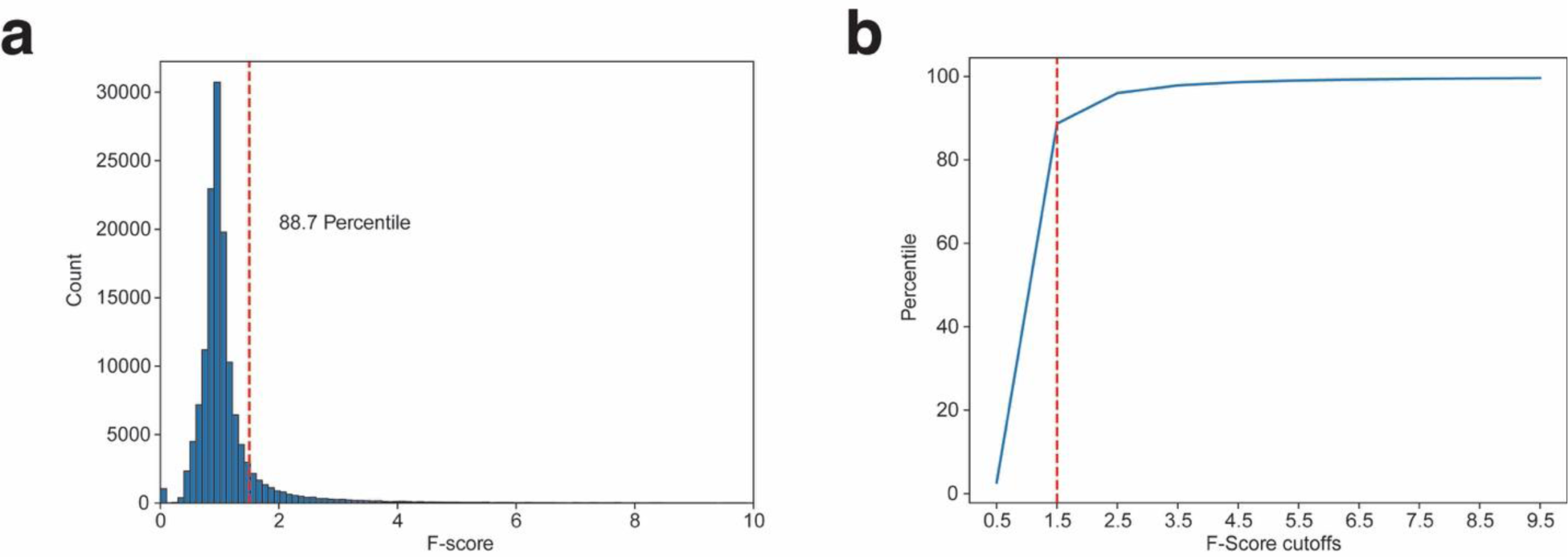
F-score threshold selection for “hits”. (a) Histogram showing all non-zero F-scores for eight ligands. The x-axis is limited to F-score of 10. Red line marks region where the F-score is 1.5. The percentile at the 1.5 F-score is noted. (b) Plot showing the percentile at a series of F-score cutoffs for the dataset containing all scores from (a). The cut-off of 1.5 represents the approximate “elbow” of the plot, which corresponds to the point of diminishing returns for higher F-score cut-offs.

**Supplementary Table 2:** Ligand concentrations

| **Ligand** | **Concentration (mol/L)** | **Solvent** |
| --- | --- | --- |
| Tamoxifen (Tam) | 0.00005 | ETOH |
| Endoxifen (End) | 0.00005 | ETOH |
| 4-hydroxy tamoxifen (4Hy) | 0.00005 | ETOH |
| N-desmethyl tamoxifen (Ndes) | 0.00005 | ETOH |
| Naltrexone (Nal) | 0.001 | Water |
| Quinine (Quin) | 0.0005 | DMSO |
| Ellagic Acid (EIIA) | 0.00015 | DMSO |
| Naringenin (Nar) | 0.001 | DMSO |
| Phloretin (Phlo) | 0.0003 | DMSO |

**Supplementary Table 3:**
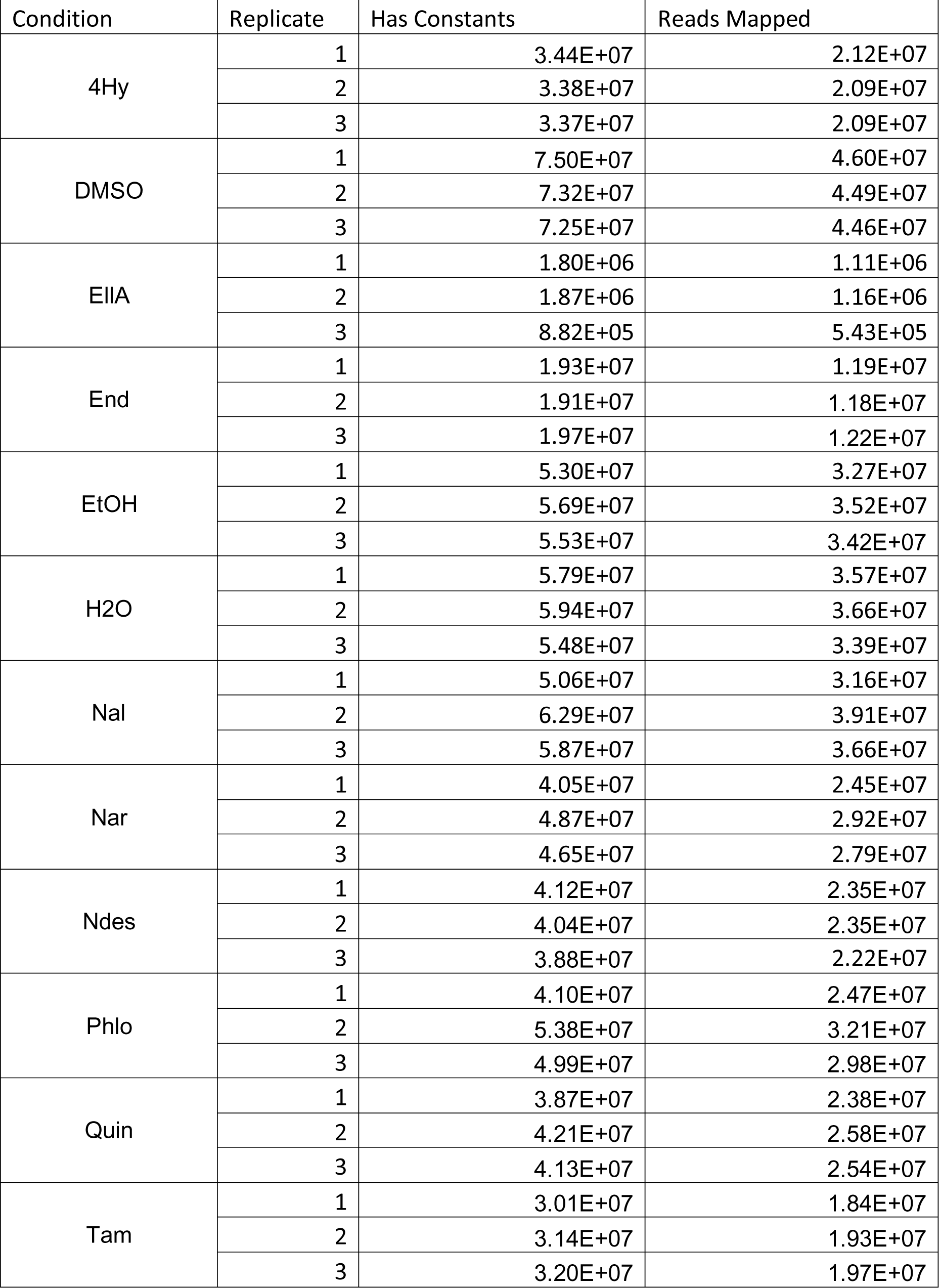
Sensor-seq DNA read counts

| Condition | Replicate | Has Constants | Reads Mapped |
| --- | --- | --- | --- |
| 4Hy | 1 | 3.44E+07 | 2.12E+07 |
|  | 2 | 3.38E+07 | 2.09E+07 |
|  | 3 | 3.37E+07 | 2.09E+07 |
| DMSO | 1 | 7.50E+07 | 4.60E+07 |
|  | 2 | 7.32E+07 | 4.49E+07 |
|  | 3 | 7.25E+07 | 4.46E+07 |
| EIIA | 1 | 1.80E+06 | 1.11E+06 |
|  | 2 | 1.87E+06 | 1.16E+06 |
|  | 3 | 8.82E+05 | 5.43E+05 |
| End | 1 | 1.93E+07 | 1.19E+07 |
|  | 2 | 1.91E+07 | 1.18E+07 |
|  | 3 | 1.97E+07 | 1.22E+07 |
| EtOH | 1 | 5.30E+07 | 3.27E+07 |
|  | 2 | 5.69E+07 | 3.52E+07 |
|  | 3 | 5.53E+07 | 3.42E+07 |
| H2O | 1 | 5.79E+07 | 3.57E+07 |
|  | 2 | 5.94E+07 | 3.66E+07 |
|  | 3 | 5.48E+07 | 3.39E+07 |
| Nal | 1 | 5.06E+07 | 3.16E+07 |
|  | 2 | 6.29E+07 | 3.91E+07 |
|  | 3 | 5.87E+07 | 3.66E+07 |
| Nar | 1 | 4.05E+07 | 2.45E+07 |
|  | 2 | 4.87E+07 | 2.92E+07 |
|  | 3 | 4.65E+07 | 2.79E+07 |
| Ndes | 1 | 4.12E+07 | 2.35E+07 |
|  | 2 | 4.04E+07 | 2.35E+07 |
|  | 3 | 3.88E+07 | 2.22E+07 |
| Phlo | 1 | 4.10E+07 | 2.47E+07 |
|  | 2 | 5.38E+07 | 3.21E+07 |
|  | 3 | 4.99E+07 | 2.98E+07 |
| Quin | 1 | 3.87E+07 | 2.38E+07 |
|  | 2 | 4.21E+07 | 2.58E+07 |
|  | 3 | 4.13E+07 | 2.54E+07 |
| Tam | 1 | 3.01E+07 | 1.84E+07 |
|  | 2 | 3.14E+07 | 1.93E+07 |
|  | 3 | 3.20E+07 | 1.97E+07 |

**Supplementary Table 4:** Sensor-seq RNA read counts

| Condition | Replicate | Has Constants | Reads Mapped |
| --- | --- | --- | --- |
| 4Hy | 1 | 2.10E+07 | 9.66E+06 |
|  | 2 | 2.22E+07 | 1.02E+07 |
|  | 3 | 2.21E+07 | 1.01E+07 |
| DMSO | 1 | 2.38E+07 | 1.04E+07 |
|  | 2 | 2.55E+07 | 1.12E+07 |
|  | 3 | 2.41E+07 | 1.05E+07 |
| EIIA | 1 | 1.55E+07 | 6.70E+06 |
|  | 2 | 2.32E+07 | 1.00E+07 |
|  | 3 | 2.32E+07 | 1.00E+07 |
| End | 1 | 2.43E+07 | 1.14E+07 |
|  | 2 | 2.33E+07 | 1.09E+07 |
|  | 3 | 2.54E+07 | 1.19E+07 |
| EtOH | 1 | 1.96E+07 | 8.70E+06 |
|  | 2 | 2.28E+07 | 1.01E+07 |
|  | 3 | 2.33E+07 | 1.03E+07 |
| H2O | 1 | 2.09E+07 | 8.98E+06 |
|  | 2 | 2.19E+07 | 9.36E+06 |
|  | 3 | 2.26E+07 | 9.77E+06 |
| Nal | 1 | 1.90E+07 | 8.31E+06 |
|  | 2 | 2.18E+07 | 9.41E+06 |
|  | 3 | 2.15E+07 | 9.35E+06 |
| Nar | 1 | 2.47E+07 | 2.47E+07 |
|  | 2 | 2.77E+07 | 1.50E+07 |
|  | 3 | 2.68E+07 | 1.45E+07 |
| Ndes | 1 | 2.28E+07 | 1.09E+07 |
|  | 2 | 2.31E+07 | 1.10E+07 |
|  | 3 | 2.44E+07 | 1.17E+07 |
| Phlo | 1 | 2.52E+07 | 1.37E+07 |
|  | 2 | 2.71E+07 | 1.47E+07 |
|  | 3 | 2.72E+07 | 1.47E+07 |
| Quin | 1 | 2.13E+07 | 9.92E+06 |
|  | 2 | 2.39E+07 | 1.11E+07 |
|  | 3 | 2.40E+07 | 1.11E+07 |
| Tam | 1 | 2.60E+07 | 1.23E+07 |
|  | 2 | 2.49E+07 | 1.17E+07 |
|  | 3 | 2.44E+07 | 1.13E+07 |

**Supplementary Table 11:** 3A7 crystallography data collection and refinement

|  | **3A7 with naltrexone** |
| --- | --- |
| **Wavelength** |  |
| **Resolution range** | 27.6 - 2.08 (2.154 - 2.08) |
| **Space group** | P 1 |
| **Unit cell** | 43.1 43.169 115.826 99.071 97.016 95.973 |
| **Total reflections** | 154547 (13986) |
| **Unique reflections** | 44735 (4030) |
| **Multiplicity** | 3.5 (3.5) |
| **Completeness (%)** | 91.56 (81.13) |
| **Mean I/sigma(I)** | 9.34 (2.35) |
| **Wilson B-factor** | 38.13 |
| **R-merge** | 0.07726 (0.4412) |
| **R-meas** | 0.0914 (0.5196) |
| **R-pim** | 0.04845 (0.273) |
| **CC1/2** | 0.994 (0.894) |
| **CC\*** | 0.998 (0.972) |
| **Reflections used in refinement** | 44675 (4020) |
| **Reflections used for R-free** | 1799 (158) |
| **R-work** | 0.2041 (0.2599) |
| **R-free** | 0.2532 (0.3148) |
| **CC(work)** | 0.961 (0.886) |
| **CC(free)** | 0.935 (0.746) |
| **Number of non-hydrogen atoms** | 7007 |
| **macromolecules** | 6641 |
| **ligands** | 292 |
| **solvent** | 212 |
| **Protein residues** | 827 |
| **RMS(bonds)** | 0.003 |
| **RMS(angles)** | 0.82 |
| **Ramachandran favored (%)** | 98.9 |
| **Ramachandran allowed (%)** | 1.1 |
| **Ramachandran outliers (%)** | 0 |
| **Rotamer outliers (%)** | 0.86 |
| **Clashscore** | 4.9 |
| **Average B-factor** | 52.38 |
| **macromolecules** | 51.93 |
| **ligands** | 74.01 |
| **solvent** | 50.85 |
| **Number of TLS groups** | 14 |
| Statistics for the highest-resolution shell are shown in parentheses. |  |

## Methods

### Plasmid creation

PCR amplicons are generated using Kapa HiFi (Roche) PCR kits following the manufacturer protocol (Supplementary Table 1). Amplicons are treated with 15U of Dpn1 (NEB) for 2.5 hours at 37°C followed by 20 minutes at 80°C. Amplicons are then purified using EZNA Cycle Pure kits (Omega BioTek). Isothermal assembly followed Gibson Assembly protocols (NEB), but contained 100 mM Tris-HCl pH 7.5, 20 mM MgCl_2_, 0.2 mM dATP, 0.2 mM dCTP, 0.2 mM dGTP, 10 mM DTT, 5% PEG-8000, 1 mM NAD+, 4 U/ml T5 exonuclease, 4 U/μl Taq DNA ligase, and 25 U/ml Phusion polymerase. Isothermal assembly reactions are diluted 10X in dH2O prior to transformation. DH10B (NEB) electrocompetent cells are transformed with 2μL of diluted isothermal assembly reaction. Transformants are recovered in 700μL SOC for 1 hour at 37°C. Dilutions are plated on LB-kanamycin (50μg/mL) plates and incubated at 37°C overnight. Colony PCR is performed using Kapa Robust (Roche) using a single colony diluted in 100μL of dH2O. Plasmid purifications are performed using the ZR Plasmid Miniprep Classic kit (Zymo). All plasmid assemblies follow this methodology unless stated otherwise.

We first generated a sensor-reporter plasmid containing a TtgR gene under an apFAB61-BBaJ61132 constitutive operator sequence, an sfGFP gene under control of the TtgR operator sequence, a kanamycin resistance cassette, and a ColE1 origin of replication^65^. The sfGFP fragment with a modified TtgR operator sequence was amplified from the TtgR_pBBR1_V2 plasmid. The TtgR gene was amplified from the TtgR_SC101BBa plasmid. The sfColE1 backbone amplicon contains a kanamycin marker and a ColE1 origin. These fragments were assembled in a Gibson Assembly reaction as described above to form TtgR_ColE1_SPS. The sfGFP promoter was then modified to have the wildtype TtgR operator sequence. sfGFP with the wildtype operator sequence was amplified from a separate plasmid. The backbone amplicon was amplified from TtgR_ColE1_SPS and consisted of the TtgR gene, the kanamycin resistance marker, and the ColE1 origin. The assembled plasmid was labeled as TtgR_ColE1_SPS_V2 and is used as the vector for cell-based fluorescent assays. A third Gibson assembly reaction was required to insert stop codons and BsaI cut sites into the middle of the GFP gene to create the barcode insertion site. The stop codons and BsaI sites were encoded on overlapping primers and added to the TtgR_ColE1_SPS_V2 plasmid. The backbone was annealed to itself in a 1-part isothermal assembly. This construct was labeled TtgR_ColE1_SPS_V5 and used as the template for the RNA-Seq assays.

### Library creation

Plasmid libraries are generated using Golden Gate Assembly Kits (NEB, BsaI-HFv2). The reactions undergo a cycling protocol of 30 alternating 5-minute 37°C and 16°C cycles followed by a final 60°C 5-minute hold. The reactions are dialyzed against dH2O on semi-permeable membranes (Millipore) for 1 hour at room temperature. DH10B (NEB) cells were transformed with 3μL of dialyzed reaction via electroporation.

Transformants were recovered in 1mL of SOC and then diluted 2X, 5X, and 10X with fresh SOC. Each dilution recovered for 1 hour shaking at 37°C. 4mL of LB-kanamycin (50μg/mL) was added to each dilution after recovery and 50X and 500X dilutions were plated of each recovered dilution to calculate transformation efficiency. The remaining transformants were grown for 6 hours shaking at 225rpm. A frozen stock was made in 25% glycerol and stored at -80°C for each dilution. Fresh cultures were created by diluting each 6-hour growth 50X into fresh LB-kanamycin. These were grown overnight, and plasmids were harvested via ZR Plasmid Miniprep Classic kit (Zymo). All plasmid library assemblies follow this methodology unless stated otherwise.

Pre-defined or random barcodes were synthesized as a short primer (IDT). These barcode primers were combined separately with another constant primer to create short double-stranded fragments containing the barcode flanked by BsaI cut sites in a single cycle of PCR using Kapa HiFi (Roche). 1μL of this reaction was added into a second Kapa HiFi (Roche) reaction with additional primers to increase the length of the amplicon over 18 cycles. The resulting amplicon was purified using the DNA Clean and Concentrator-5 kit (Zymo).

To generate the pilot 16-member mini-library, the TtgR gene variants were isolated from a set of 16 pre-existing plasmids each containing a single TtgR variant^17^. 100ng of each amplicon was combined in a pool. The TtgR_ColE1_SPS_V5 backbone was amplified using primers that encompassed the sfGFP gene, the ColE1 origin, and the kanamycin resistance marker. The barcodes, TtgR gene variants, and backbone were assembled in a single Golden Gate reaction (NEB) as described above.

FuncLib mutations were encoded into short oligos (Agilent) consisting of the TtgR gene region flanked by BsaI cut sites for Golden Gate assembly. Four pools of approximately 4,400 variants were created by randomly combining between 1 and 4 tolerated mutations. Each pool had unique priming sequences to isolate from a pooled sample.

The pooled library was diluted to 0.005μM in Tris-HCl (pH 7.5). Each pool was amplified using Kapa HiFi and 1μL of the diluted library in 15 cycles in triplicate. The amplified reactions were pooled together and purified using the DNA Clean and Concentrator 5 kit (Zymo). The pooled oligos, barcodes, and TtgR_ColE1_SPS_V5 backbone were assembled using Golden Gate assembly (NEB). The libraries with approximately 15 barcodes per variant, calculated by CFU/mL, were selected for RNA-Seq.

### Creating GFP control

To create a GFP positive control, the TtgR gene was removed from the TtgR_ColE1_SPS_V2 plasmid. The backbone was amplified with primers that had complementary overlap with the sfGFP gene. The sfGFP gene was amplified with primers complementary to the backbone. The BsaI cut sites and early stop codons were inserted into sfGFP in the same fashion as the creation of TtgR_ColE1_SPS_V5. The plasmid was labeled TtgR_ColE1_SPS_V3_GFPControl. Three pre-defined 20nt barcodes (AAACCCTGTGCCAGAGGGTG, GAGTGACCTTAAGTCAGGGA, and GCTTCTGTCCAAGCAGGTTA) were generated according to standard protocols. The barcodes were inserted into the TtgR_ColE1_SPS_V3_GFPControl using Golden Gate assembly.

### RNA preparation for Sensor-Seq

For the pilot study, cells containing 8 different TtgR variants were streaked out on an LB-Kan plate and grown overnight at 37°C. Three colonies were inoculated into LB-kanamycin for overnight growth. The overnight cultures were diluted 50X into fresh LB-kanamycin containing either1mM naringenin or DMSO as a control. DH10B containing each barcoded GFP Control plasmid were struck out on LB-kanamycin plates. One colony was selected from each barcoded DH10B and grown in 3mL LB-kanamycin overnight. These barcoded control cultures were combined in equal ratio and added to the test library culture to a final composition of 0.25% control. The induced cultures were grown and prepared following standard protocols The cultures were grown at 37°C shaking at 250rpm in an Innova 4230 (New Brunswick Scientific). At the targeted OD600, cultures were placed on ice for 10 minutes. 5E8 cells were harvested by centrifugation at 5,500g based on the OD600 and the assumption that 1.0 OD600 cultures have 8E8 cells/mL. The pelleted cells were decanted and stored at -80°C. This process was repeated in biological triplicate for each target OD600 with new colonies.

RNA was purified from cell pellets via Trizol reagent (Invitrogen). 1mL of Trizol reagent (Invitrogen) was added to each cell pellet and vortexed briefly. The samples incubated at room temperature for 5 minutes. 200μL of chloroform (Sigma Aldrich) was added to each sample. The samples incubated at room temperature for 2 minutes and were centrifuged at 12,000g for 15 minutes at 4°C. 300μL of the aqueous phase was transferred to a clean 2mL centrifuge tube and placed on ice. RNA was purified from the aqueous phase using the RNA Clean and Concentrator 5 kit (Zymo) and eluted in 15μL Ultrapure RNase-free dH2O (Invitrogen). The purified RNA was digested using 4U DnaseI (NEB) in a 50μL reaction incubated at 37°C for 30 minutes. The digestion reactions were purified using the RNA Clean and Concentrator 5 kit (Zymo) and eluted in 15μL Ultrapure RNase-free dH2O (Invitrogen). Concentrations were measured using a Nanodrop instrument (Thermo Fisher).

To test the agnostic library, the four pools were grown individually in 5mL LB-kanamycin overnight in triplicate. The four pools were combined prior to inoculation in 25mL LB-kanamycin for the RNA harvest. GFP Control barcoded cells were spiked into the combined agnostic replicates at a final concentration of 0.25%. The same pooled replicates were used for all ligand inductions. The agnostic libraries were induced with the ligand (Supplementary Table 2). DMSO, dH2O, and EtOH were included as solvent controls. No more than 2% v/v (DMSO) or 1% v/v (EtOH and H_2_O) of solvent were tolerated. These cultures were processed as previously done with the pilot library.

### cDNA synthesis and sequencing

cDNA synthesis uses approximately 3μg total RNA, a primer encoding a 16nt unique molecular identifier (UMI), and the Maxima H Minus Double-Stranded cDNA Synthesis Kit. The cDNA is purified using the DNA Clean and Concentrator 5 kit (Zymo). The Illumina sequencing regions are added in 2 PCR reactions in the same manner as the MiSeq barcode-variant mapping reactions. Three sets of primers containing the Illumina sequencing primer and a predefined barcode (ATCG, CGAT, and GTCA) were used in the first PCR reaction to add the Illumina sequencing regions (11 cycles). One set of primers was used for each biological replicate. The first reaction is purified using the DNA Clean and Concentrator 5 kit (Zymo). The second reaction uses 4μL of the first reaction and primers that add i5 and i7 indices in 8 cycles. The final amplicons are purified again. All replicates were combined in an equal molar ratio after purification.

Plasmids are harvested from the remaining culture of the RNA preparation step. The UMI is added to the plasmid-derived samples in a 2-cycle PCR reaction using 100ng of template. The amplification of all DNA libraries followed an identical protocol to the RNA preparation.The cDNA and DNA samples are sequenced using either a NovaSeq SP chip (test library) or a NovaSeq S4 chip (agnostic libraries) by the UWBC. Read volumes were calculated by targeting 500 reads per barcode with the assumption that 50% of the reads will be lost due to filtering criteria.

### RNA-Seq Data Analysis

Fastq files were merged using NGmerge and filtered using Fastp based on average Q-score > Q30 for reads^66,67^. Reads containing the 5’ and 3’ constant regions were isolated using UMI-Tools and counted using Tally^68,69^. Reads containing the central constant region were isolated and UMI sequences were removed with UMI-Tools. The barcodes were then counted with Tally. RNA-Seq barcodes were matched to mapped barcode-variant pairs with a Hamming distance tolerance of 1 using Seal (sourceforge.net/projects/bbmap/). If a barcode mapped to more than one TtgR variant, then the TtgR variant that had the most reads was selected if each other variant was less than 10% of the reads of the most abundant variant. RNA-Seq barcodes that were successfully mapped to known barcode-variant pairs were analyzed across the induced RNA, induced DNA, control RNA, and control DNA samples. A barcode both had to be found in all four groups to be included in downstream analysis. The read counts for a variant were then a sum of the barcode counts for all barcodes mapped and found in all four datasets. No read count threshold was imposed during analysis. The fold enrichment calculation uses equation (1).

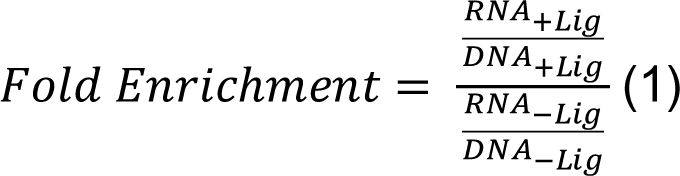

If biological replicates were available for each condition, the fold enrichment per variant was curated based on the coefficient of variation (CV). Percent deviation is calculated with equation (2).

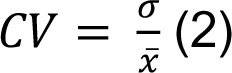

In this equation, σ is the standard deviation of the fold enrichment and ̅𝑥 is the mean fold enrichment across replicates. A 30% CV cutoff was imposed (Supplementary Fig. 3). All variants were normalized to wildtype fold enrichment for each replicate.

Heatmaps were constructed using the average performance of each variant after normalization. Variants with data passing CV thresholds for more than 5 ligands and performed at least 1.5 times better than wildtype were selected for clustering. Missing data was imputed using KNN methods in SciKit Learn^70^. The UPGMA algorithm with a correlation distance metric and a target of 12 clusters was used to cluster in SciPy^71^. The number of clusters was selected by plotting the silhouette score against the number of clusters (Supplementary Fig. 5).

### Mapping barcode-variant pairs

A 60nt spacer was created to bring the random barcode and TtgR variants physically adjacent on the same plasmid to enable short read next generation sequencing mapping of barcode-variant pairs. The library plasmids were amplified with primers encoding BsaI cut sites that would place the spacer between the barcode and the TtgR variant region. The spacer was inserted into the backbone using Golden Gate (NEB) following standard protocols.

Two primer groups were used to add Illumina sequencing regions to the barcode-spacer-variant region of the mapping plasmid libraries. Each primer group consisted of three primers with different numbers of Ns (0N, 3N, or 6N) to increase positional base diversity during runs. The adapter primers had complementarity to the plasmid and contained Illumina sequencing primer binding regions. Stem primers had the i7 and i5 indices and the adapter sequence to anneal to the sequencing flow cell. The adapter regions were added using 1ng of template, 0.6μL of 10μM primers, and Kapa HiFi mix (Roche) for 14 cycles. These reactions were purified using the DNA Clean and Concentrator 5 kit (Zymo). The stem primers were used in a second PCR reaction using 4μL of the first reaction for 10 cycles.

The pilot library was sequenced on a 15M 2×250 MiSeq chip (Illumina). The agnostic library was sequenced using an 2×250 NovaSeq SP chip (Illumina). For MiSeq-based sequencing, the proper band was isolated using gel extraction on a 0.5% agarose gel followed by purification with the EZNA gel extraction kit (Omega BioTek). The concentration of the DNA was measured using AccuClear (Biotium) following manufacturer protocols. The flow cell was loaded with 15pM DNA with 5% PhiX. For NovaSeq-based sequencing, samples were purified using PippinHT (Sage Science) and the concentration was measured via 4200 TapeStation (Agilent). The size selection, concentration measurement, and NovaSeq runs were performed by the University of Wisconsin Madison Biotechnology Center (UWBC).

The fastq output was merged using PEAR^72^. A C++ script was used to filter poor-scoring reads based on Q-scores. Reads that passed the quality filter were then filtered on constant regions surrounding the barcode and TtgR variants. Barcodes that had read counts greater than 10 and were unique for a single TtgR variant were mapped to that variant. If a barcode mapped to more than one TtgR variant, then the TtgR variant that had the most reads was selected if each other variant was less than 10% of the reads of the most abundant variant.

### qRT-PCR quantification of transcript abundance

The abundance of the sfGFP and rrsA transcripts were measured via qRT-PCR. Each biological triplicate RNA was run in a technical triplicate in a MicroAmp Fast Optical 96-well plate (Life Technologies). 1ng of RNA was added to Luna Universal One-Step qRT-PCR mix (NEB) containing 4μmol of each primer on ice. The standard cycling protocol was used according to the manufacturer’s suggestion. Each sample consisted of a set of reactions containing sfGFP-specific primers and another set containing rrsA-specific primers. The reactions were run on a CFX Connect Real Time PCR Detection System (BioRad). Fold enrichment was calculated using equations (3) and (4). The error was propagated from the technical replicates and biological replicates using (5).

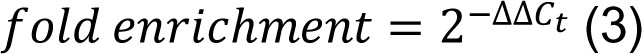

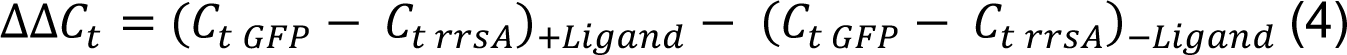

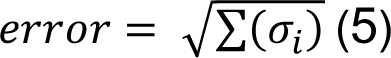

### Cell Sorting

An overnight culture is diluted 50X in phosphate buffered saline (137mM NaCl, 2.7mM KCl, 10mM Na2HPO4, 1.8mM KH2PO4) and placed on ice for 10 minutes prior to sorting. Sorting was performed on an SH800 (Sony) using the 488nm laser and a 525±25nm filter. Sorted cells were grown for 1 hour shaking at 37°C in 5mL LB. Kanamycin was added to a final concentration of 50μg/mL and the culture was grown overnight. An aliquot of the sorted culture was stored at - 80°C in 25% glycerol. Plasmids were isolated from the remaining culture using the ZR Plasmid Miniprep – Classic kit (Zymo).

### Secondary Validation of Top Hits

Top performing variants were selected based on the mean rank of each variant across the three biological replicates. These variants were encoded in gene fragments (Twist) and synthesized in a 96-well plate format. The fragments were resuspended to a final concentration of 10ng/μL, pooled together, and cloned into the TtgR_ColE1_SPS_V2 backbone using Golden Gate Assembly. The resulting library was sorted based on fluorescence. LB-kanamycin is inoculated with 50μL of the frozen stock of the library and grown overnight shaking at 37°C. Sorting was performed according to the Cell Sorting protocol. 500,000 cells were isolated from the lower 70% of the population based on fluorescence. Plasmids were isolated from the remaining culture using the ZR Plasmid Miniprep – Classic kit (Zymo). DH10B (NEB) were transformed with the purified plasmid library according to the Library Creation protocol.

LB-kanamycin is inoculated with 50μL of the frozen stock of the repressed library and grown overnight, shaking at 37°C. The culture was diluted 50X into fresh LB-kanamycin and grown overnight at 37°C shaking with the ligands (Supplementary Table 2). Sorting was performed according to the Cell Sorting protocol. 400,000 cells were isolated using a gate that encompassed the top 0.5% of the population based on the fluorescence distribution in the absence of any ligand. Plasmids were isolated from the remaining culture using the ZR Plasmid Miniprep – Classic kit (Zymo).

The abundance of variants was determined using next-generation sequencing. Sequencing amplicons were generated using primers that had complementarity to the TtgR gene around the gene fragment insertion site. The amplification process followed the “Barcode-variant mapping via next-generation sequencing” protocol. The concentration of the DNA was measured using Qubit Fluorometric Quantification (Thermo Fisher) following manufacturer protocols. The flow cell was loaded with 15pM DNA with 5% PhiX. Sequencing was performed on a MiSeq instrument (Illumina). Fastq files were merged using NGmerge and filtered using Fastp based on average Q-score > Q30 for reads^42,43^. The abundances of each variant in each sample, including the repressed population, were normalized to the total reads for that sample. The fold enrichment is the ratio of the induced and repressed normalized abundances.

### Tertiary Validation of Top Hits

Top three performing variants for each ligand were selected based on the secondary validation. If a top variant for a ligand exists in the current selection, the next best variant is selected until each ligand has three top performers. These variants are individually cloned into the TtgR_ColE1_SPS_V2 backbone using Golden Gate Assembly. 3uL of overnight culture for each clone is inoculated into 144uL LB-kanamycin spiked with 3uL of ligand or vehicle control in triplicate such that the final working concentration is as described in Supplementary Table 2. These cultures were grown at 37°C shaking overnight. Induced overnight cultures were diluted 50X in 1X PBS. 10^2^-10^3^ cells were collected using an Attune NxT Flow Cytometer with Autosampler (Thermo Fisher). The median fluorescence of the collected cells for each sample was determined. The fold induction is calculated as the median fluorescence in the presence of ligand divided by the median fluorescence in the absence of ligand.

### Unsupervised learning of ligand-agnostic dataset

The 11 possible mutated positions of the TtgR agnostic library were each physicochemically encoded in a 19-length dimenstionally-reduced AAindex for a total of 209 features for each variant^44,45^. The variants were embedded in a 2-dimensional projection and subsequently clustered using a combination of UMAP and HDBSCAN^43,47^. Hyperparameter space (n_neighbors: [15, 25, 50, 75, 100, 300, 500, 1000, 2000], n_components: [2], min_dist: [0, 0.1], metric: [manhattan], min_cluster_size: range(5, 110, step=5)) was randomly searched with 100 iterations to find the optimal hyperparameters that minimized a cost function. Here, the cost function is the percentage of points with <5% certainty of assignment. The best combination was found to be n_neighbors=500, min_dist=0.1, and min_cluster_size=80, which minimized the cost function to 0.03144 (i.e. ∼3% of points had <5% certainty of assignment). These hyperparameters were used to cluster the points in the associated UMAP embedding.

Top three clusters for each ligand were chosen for further analysis. Cluster rankings were determined by the average F-score of “hits” (variants with F-score ≥ 1.5). In order to be considered as a top three cluster, we also require the cluster to have a minimum of 15 “hits”. To determine the enrichment of substitutions at each mutated position from these top clusters, we calculated the ratio of the F-score-weighted substitution frequency within “hits” with the DNA-count-normalized substitution frequency. The F-score-weighted substitution frequency is determined by finding the frequency of a substitution in a set of variants where each variant’s abundance is weighted by its average F-score from three replicates. The DNA-count-normalized substitution frequency measures the average frequency of each substitution in the induced plasmid library where the abundance of each variant is dictated by the number of sequenced DNA barcodes associated with that variant and is used to control for bias that may exist in the plasmid library. Thus, the ratio of these two values describes the change in frequency of each substitution such that large values signify enrichment of a substitution in presence of a ligand.

### Cell-Free Extract Preparation

Cell-free extract was prepared as previously described with some modifications^54^. 1 L of 2X YT + P media (16 g/L tryptone, 10 g/L yeast extract, 5 g/L NaCl, 7 g/L potassium phosphate dibasic, 3 g/L potassium phosphate monobasic) was inoculated with 20 mL of saturated overnight culture of BL21 Star (DE3) in LB and grown to optical density 2.6. The cells were harvested by centrifugation at 5,000 X g for 10 minutes, resuspended and washed in Buffer B (14 mM Mg-glutamate, 60 mM K-glutamate, 5 mM Tris, pH 8.2) three times, then resuspended to a final concentration of 1 g/mL cell in Buffer B. The suspension was lysed using a QSonica Q125 sonicator with a 3.175 mm diameter probe at a frequency of 20 kHz and 50% amplitude by 10 second ON/OFF pulses until the lysed suspensions turned brown and became less viscous (around 60 seconds and delivering ∼ 350 J). The lysate was clarified by a 10-minute centrifugation at 12,000 X g and 4 °C. The supernatant was removed and incubated, shaking at 220 rpm for 80 minutes at 37 °C for the ribosomal runoff reaction. After a second 12,000 X g spin at 4°C for 10 minutes, the supernatant from the runoff was dialyzed against Buffer B overnight at 4 °C in a 3.5K MWCO membrane. The dialysate was removed, centrifuged once more, aliquoted, and flash-frozen on liquid nitrogen for long-term storage at -80°C.

### Cell-Free Gene Expression Reaction

CFE reactions were prepared as previously described^54^. The overall reaction composition was 6 mM magnesium glutamate; 10 mM ammonium glutamate; 130 mM potassium glutamate; 1.2 mM ATP; 0.850 mM each of GTP, UTP, and CTP; 0.034 mg/mL folinic acid; 0.171 mg/mL yeast tRNA; 2 mM amino acids; 30 mM PEP; 0.33 mM NAD; 0.27 mM CoA; 4 mM oxalic acid; 1 mM putrescine; 1.5 mM spermidine; 57 mM HEPES; 30% extract by volume; plasmid DNA to the desired concentration and water. 10 µl reactions were mixed on ice in replicates and then pipetted onto a black Corning clear-bottom 384-well plate for measurement of sfGFP (excitation/emission 470/510 nm) on a Biotek Synergy H1 plate reader every ten minutes at 30 °C.

### X-ray Crystallography

TtgR-pET31B variants were expressed in BL21 cells (NEB) as previously described with a few minor differences^33^. 1 L of TB with 20xM and 80155 autoinduction media^73^ was inoculated from a starter culture and grown at 25°C for 24 h. The cells were harvested by centrifuging 20 min at 5000 rpm. To purify, cells were lysed using an M110 Microfluidizer (Microfluidics International Corporation), and purification was carried out as previously described^33^.

Crystals were screened based on previously published conditions^33^, however, only variant 3A7 yielded diffraction quality crystals in 100 mM Bis-Tris pH 6.5 18-20% MEPEG 2000, 200 mM MgSO_4_ after extensive effort. To obtain naltrexone in the active site, crystals were both co-crystalized with 5 mM naltrexone and prior to cryoprotection, were incubated for 1 h in cryoprotectant (100 mM Bis-Tris pH 6.5 18-20% MEPEG 2000, 200 mM MgSO_4_ 20% glycerol) with saturating naltrexone and frozen in lN_2_.

Xray diffraction data were collected at Advanced Photon Source (APS) beamlines LS-CAT ID-D. Diffraction data was reduced and scaled using XDS and autoPROC^74,75^. The structure was solved by molecular replacement with Phenix.phaser^76^ using PDB ID 7K1C^33^. The model was manually inspected and built through iterative rounds of Phenix.refine and manual inspection in COOT^77^.

## Supporting information

supptables_fig3

supptables_fig2

## Acknowledgements

This was supported by United States Army Research Office Grants W911NF20C0005 and W911NF1710043 (S.R.). K.K.N was supported by NIH National Research Service Award T32 GM07215 (Molecular Biophysics Training Program) and the Robert and Katherine Burris Biochemistry Fund. J.C was supported by the National Institute of General Medical Sciences of the National Institute of Health under Award Number T32GM135066 (Biotechnology Training Program). S.H and J.L.C were supported by the AFRL 711th Human Performance Wing.

## Notes

### Competing Interest Statement

The authors have filed a provisional patent application for this technology

### Summary of Updates

Updated abstract, main Figure 2, and text from Introduction up to text corresponding to main Figure 3.

